# Cornichon receptors couple membrane adaptation to cargo selection during ER export

**DOI:** 10.64898/2026.08.07.743479

**Authors:** Jude Tunyi, Oliver Adams, Siân Holton, Martyna Biaduń, Nathan Bernhardt, Gabriel Kuteyi, Omar Pantoja, Lucy R. Forrest, Joanne L. Parker, Simon Newstead

## Abstract

Selective export of membrane proteins from the endoplasmic reticulum (ER) is fundamental for eukaryotic cell biology, yet how trafficking receptors coordinate cargo recognition with membrane adaptation and COPII recruitment remains unknown. Cornichon homolog (CNIH) proteins comprise a conserved family of trafficking receptors that mediate ER export of ion channels, G protein-coupled receptors (GPCRs), ATP-binding cassette (ABC) and solute carrier (SLC) transporters. Here, we determine cryo-electron microscopy structures of the prototypical cornichon receptor Erv14 bound to an SLC transporter in detergent and lipid nanodiscs. We show that cargo recognition is mediated by a dynamic network of interactions, in which structural lipids stabilize the receptor-cargo interface. Nanodisc structures reveal the assembly of a second Erv14 receptor that remodels the receptor-cargo interface in response to membrane architecture, thereby reducing local membrane thickness and providing direct structural evidence that cornichon receptors buffer hydrophobic mismatch during membrane protein biogenesis. Structural and trafficking analyses further show that the second receptor recruits the COPII adaptor Sec24, coupling membrane remodelling to cargo export. Together, our findings establish that cornichon receptors couple lipid-mediated membrane adaptation with cargo selection through sequential receptor assembly, linking membrane protein folding to selective COPII-mediated ER export.

**One sentence summary:** Cornichon receptors integrate membrane adaptation with cargo recognition to coordinate membrane protein quality control and selective ER export.

## Introduction

Cargo selection at ER exit sites is a fundamental step in membrane protein biogenesis and quality control (*1*). Defects in membrane protein folding and trafficking underlie numerous human diseases, including cystic fibrosis, channelopathies and neurodegenerative disorders (*2–7*). Membrane proteins destined for the Golgi and plasma membrane are exported from the ER in COPII-coated vesicles (*1, 8*). Although some membrane proteins interact directly with the Sec24 component of the COPII coat complex (*9, 10*), approximately 30% of the plasma membrane proteome, including GPCRs, ion channels, ABC and SLC transporters, require dedicated trafficking receptors for ER export (*11, 12*). These cargoes differ substantially in topology, oligomeric state, and transmembrane helix architecture, posing a fundamental question in cell biology (*13*): how do trafficking receptors recognise structurally diverse membrane proteins while maintaining stringent ER export specificity?

Trafficking receptors couple membrane proteins to the COPII machinery and therefore occupy a pivotal position at the interface of protein folding, quality control and intracellular trafficking (*8, 14*). Beyond acting as cargo adaptors, several trafficking receptors have been proposed to function as folding chaperones, facilitating the insertion and maturation of membrane proteins with unusually long transmembrane helices within the relatively thin ER membrane (*15, 16*). However, direct structural evidence supporting this model has been lacking. Because cargo recognition occurs within the membrane, the surrounding lipid environment is also expected to influence receptor-cargo interactions (*17–19*), yet the contribution of membrane lipids to selective ER export remains enigmatic. A central challenge is to explain how trafficking receptors recognise membrane protein cargoes, distinguish mature export-competent proteins from folding intermediates, and accommodate the physical constraints imposed by the membrane (*10, 13*).

Cornichon (CNIH) proteins form an evolutionarily conserved family of trafficking receptors that mediate the ER export of diverse membrane proteins across fungi, plants, and animals (*15, 20–30*). Recent studies of mammalian CNIH proteins bound to AMPA receptors have established how cornichons function as auxiliary subunits that regulate channel gating (*31–37*). The conserved architecture of cornichon proteins suggests that common structural principles govern their diverse functions as auxiliary subunits and ER trafficking receptors. However, it remains unknown how this conserved family recognises structurally diverse membrane protein cargoes during ER-to-Golgi trafficking in the secretory pathway. In particular, it is unclear whether cargo recognition relies on a conserved protein-protein interface or on adaptable interaction networks that accommodate structurally distinct membrane proteins. Likewise, the mechanisms by which cornichon receptors coordinate cargo recognition with membrane adaptation, quality control and COPII recruitment remain unresolved (*10, 38–40*).

Here, we combine cryo-electron microscopy with biochemical, biophysical, and cell-biological approaches to determine the structure of the prototypical cornichon receptor, Erv14, bound to a multipass SLC transporter and show that cargo recognition is mediated by a reconfigurable network of protein-protein and protein-lipid interactions. We propose that cornichon receptors perform sequential functions during membrane protein biogenesis, in which one receptor stabilizes cargo during biogenesis whereas recruitment of a second receptor remodels the membrane and promotes COPII-driven ER export.

### Structural basis of cargo recognition

To elucidate structural details of cornichon cargo interactions, we determined the cryo-EM structure of the yeast cornichon receptor Erv14 in complex with the multipass SLC transporter Qdr2, a fungal multidrug resistance transporter, at 2.83 Å resolution (Fig. 1A, Table S1 & Fig. S1). The Erv14 receptor contains four transmembrane (TM) helices that form a compact, highly asymmetric structure with an overall topology similar to that of the CNIH1, CNIH2, and CNIH3 receptors bound to AMPA receptor subtypes (*31–37*) (Fig. 1B & Fig. S2A-D). The N- and C-termini of the longest helices, TM1 and TM4, which measure 46 and 57 Å in length, respectively, project into the ER lumen and pack together against TM2 and TM3, which measure 45 and 29 Å. The receptor is wedge-shaped due to a prominent kink in TM3 at the conserved proline P76 (Fig. S2A), which induces a 33° rotation of the cytoplasmic half of TM3 away from the membrane normal, while TM4 forms a distinctive crescent that bows outward into the membrane.

**Fig. 1.**
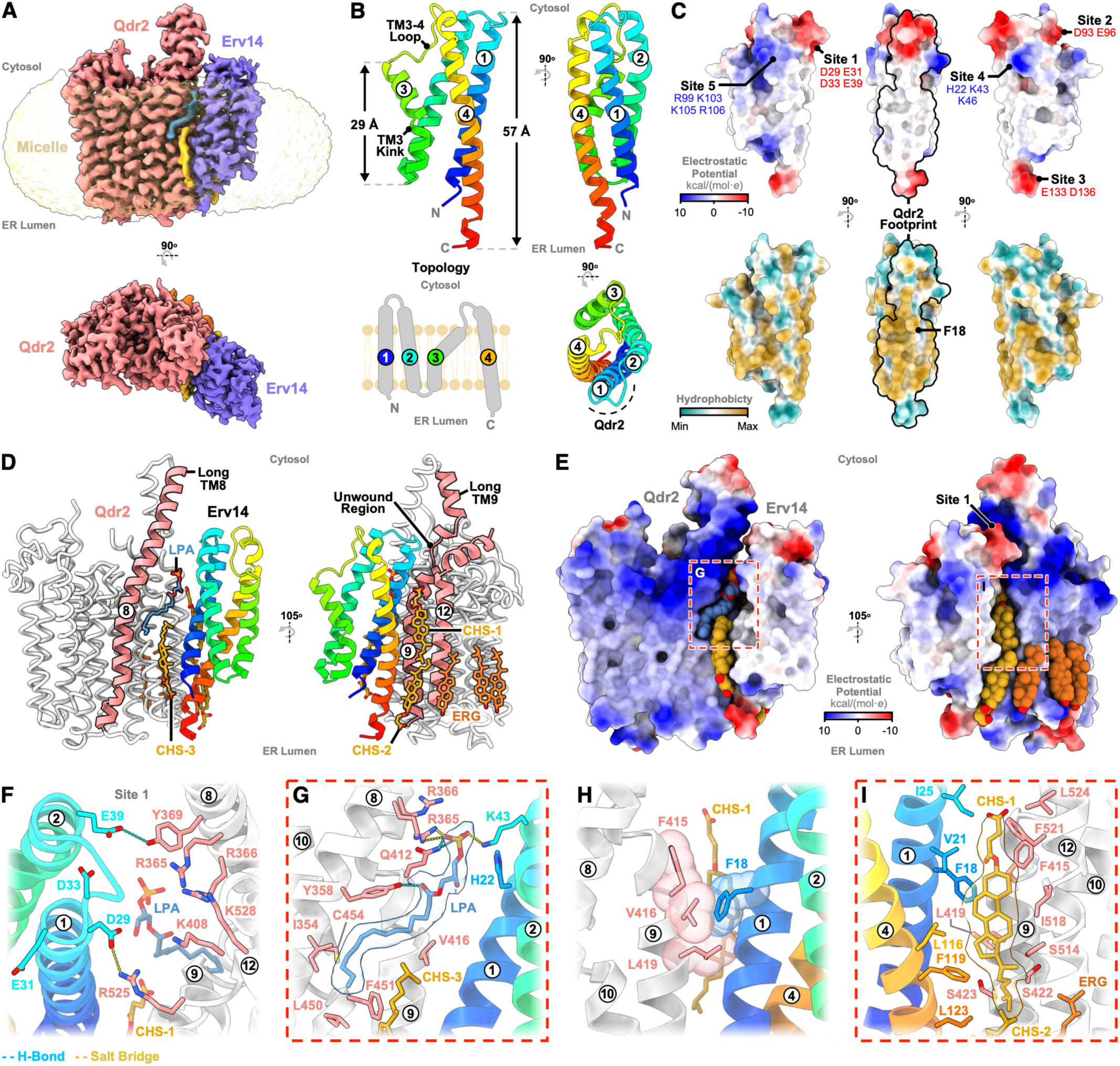
Cryo-EM structure of the yeast cornichon receptor Erv14 bound to Qdr2. **(A)** Sharpened cryo-EM map of the Erv14-Qdr2 complex (contour level 0.4), overlayed with density corresponding to the LMNG:CHS micelle (contour level 0.125, unsharpened map), with LPA (blue) and cholesterol hemisuccinate (gold) lipids. **(B)** Cartoon representation of Erv14, coloured from N-terminus (blue) to C-terminus (red), detailing the membrane topology. The curved dashed line (black) illustrates the binding interface with Qdr2 and helices 1-4 are labelled. **(C)** Surface representations of Erv14 highlighting electrostatic and hydrophobic properties; key sites are labelled. **(D)** Overview of the Erv14-Qdr2 structure. Erv14 is coloured rainbow, and Qdr2 grey in liquorice representation. All modelled lipids (LPA, CHS1-3, ERG) are shown as sticks. The elongated TM8 of Qdr2 and the unwound region of TM9, both highlighted in pink alongside TM12, are positioned adjacent to the bound interfacial lipids. **(E)** Electrostatic surface representations with interfacial lipids shown as spheres. Boxed regions correspond to the lipid interactions shown in panels G & I. Site 1 is annotated on the cytosolic surface. **(F-G)** Interaction close-ups of the Erv14-Qdr2 interface coloured as in panel D. Where shown, cryo-EM lipid density is contoured at 0.5 (sharpened map). Panels detail: **(F)** Direct protein-protein interactions at Site 1, **(G)** Coordination at the LPA binding site and **(H, I)** Hydrophobic packing of F18 of Erv14, CHS1 and Qdr2 within the membrane.

The receptor contains several conserved charged regions. Specifically, three negatively charged regions comprise D29, E31, D33, Y34 and E39, hereafter referred to as Site 1 on the TM1-TM2 cytoplasmic loop; D93 and E96, Site 2, in the unstructured region connecting TM3 and TM4; and E133 and D136, Site 3, on the luminal side of the receptor at the C-terminus of TM4. These are complemented by two positively charged regions on the receptor’s cytoplasmic side, comprising H22, K43, K46, Site 4, and R99, K103, K105, R106, Site 5 (Fig. 1C). The remainder of the interface is predominantly hydrophobic and packs against Qdr2 via TM1 and TM4, which interact with TM8, TM9, and TM12, forming a buried surface area of ∼733 Å^2^ (Table S2). Qdr2 is a member of the major facilitator superfamily (MFS) of secondary active transporters (*41, 42*), the largest family among the ∼400 SLC genes in the human genome (*43*). Although Qdr2 adopts a canonical cytoplasmic open state for an MFS protein (*44*), TM8 and TM9 are notably extended, measuring 75 Å in length (Fig. 1D), nearly double the length of canonical TM helices found in many MFS members, which are normally ∼40 Å (*45*), supporting previous models for cargo recognition by Erv14 (*15*). The cytoplasmic extensions of TM8 and TM9 contain several basic side chains, which give this region of the transporter a prominent positive charge, projecting towards Site 1 on the Erv14 receptor, where a direct salt bridge forms between D29 and R525 on TM12 of Qdr2 (Fig. 1E, F).

A notable feature of the interface is the presence of several lipid molecules. A lysophosphatidic acid (LPA) is clearly resolved in the EM volume. The phosphate head group interacts with R365 on TM8 of Qdr2 and with K43 on TM2 of Erv14 via salt bridge interactions and is positioned near H22. Additional interactions are mediated by Q412 and Y358 on Qdr2, which form hydrogen bonds with the head group and the carbonyl group of LPA, respectively, while its fatty acid chain extends back to occupy a hydrophobic cavity between TM8 and TM9 in the transporter (Fig. 1G). Three cholesterol hemisuccinate (CHS1-3) molecules are also well resolved, making direct interactions with hydrophobic residues in both Erv14 and Qdr2 and stabilising the complex (Fig. 1D, E & Fig. S2E). Three ergosterol lipids are also observed packing against the luminal side of Qdr2; however, these do not play a prominent role in the interaction with Erv14 (Fig. 1D, E). Finally, we noticed that a conserved aromatic side chain, F18 on TM1 of Erv14 (Fig. S2A), is the most prominent close contact with the Qdr2 cargo, packing against F415, V416 and L419 in TM9 in a classic ‘knobs-in-holes’ arrangement within the membrane bilayer (Fig. 1H) (*46*). Notably, this feature is also shared with CNIH receptors bound to AMPA receptors (*32, 34*). The F18 interaction of Erv14 with Qdr2 appears to be further stabilised by one of the CHS lipids (CHS1), which coordinates TM4 of the receptor with TM12 of the transporter (Fig. 1I).

### Lipids modulate the cornichon-cargo interaction

The presence of LPA and CHS at the Erv14-Qdr2 interface suggested that anionic lipids modulate Erv14-cargo interactions. Consistent with this hypothesis, CHS enhanced the association between Erv14 and Qdr2 (Fig. 2A & Fig. S2E). In the structure, the negatively charged headgroups of LPA and CHS1 are positioned between positively charged surfaces on Qdr2 (R365 and R366) and Site 4 of Erv14 (Fig. 1E, G), suggesting that they stabilise the interface through electrostatic interactions. Molecular dynamics simulations of the Erv14-Qdr2 complex in a mixture of anionic and zwitterionic lipids using a coarse-grained representation (Fig. S3A) revealed that anionic lipids (POPG) are highly enriched over the zwitterionic lipid (POPE) molecules (Fig. S3B) and localized to this charged region with long dwell times (Fig. S3C). Supporting an important role for electrostatics in the interface between the receptor and cargo, elevated ionic strength destabilised the complex (Fig. 2A), whereas 1-palmitoyl-2-oleoyl-*sn*-glycero-3-phosphate (POPA), which has a similarly small anionic headgroup, enhanced complex formation (Fig. 2B & Fig. S2E-G). Increasing phospholipid headgroup size (POPA > POPS) or reducing headgroup charge (POPA > POPC) progressively weakened the complex, indicating that receptor-cargo interactions are modulated by lipid composition. The presence of LPA and CHS at the Erv14-Qdr2 interface may help explain cholesterol enrichment in COPII vesicles (*19*) and indicates that cornichon receptors facilitate the accumulation of curvature-promoting lysophospholipids at ER exit sites (*47*).

**Fig. 2.**
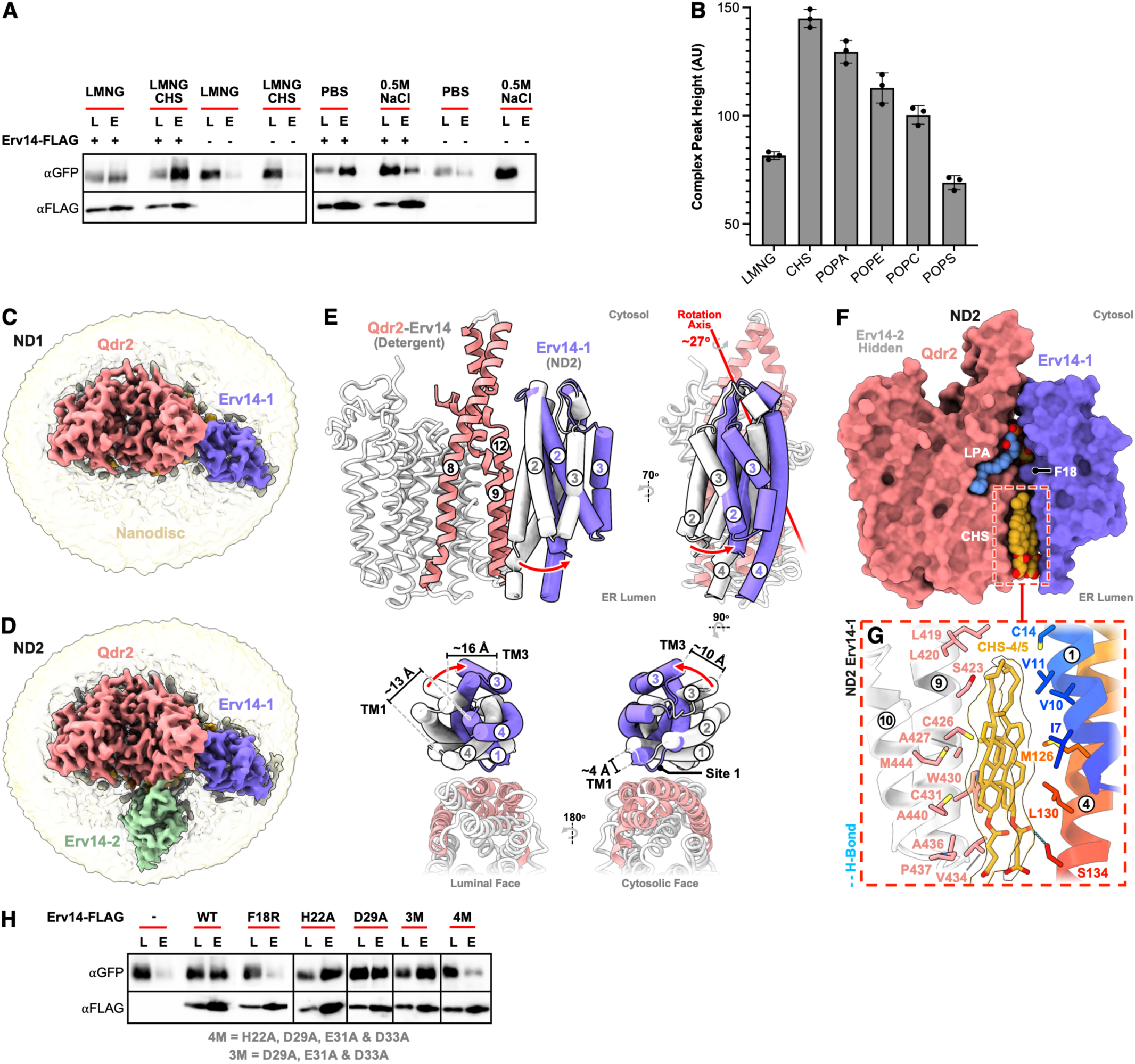
Lipids stabilize and remodel the receptor-cargo interface. **(A)** Co-immunoprecipitation of FLAG-tagged Erv14 with GFP-tagged Qdr2 under the indicated detergent (LMNG) and buffer conditions. CHS enhances complex formation, whereas elevated ionic strength disrupts the interaction. L = Load; E = Elution. **(B)** Effect of different lipids on complex formation, measured using size-exclusion chromatography peak height, in the presence of CHS, phosphatidic acid (POPA), phosphatidyl ethanolamine (POPE), phosphatidyl choline (POPC) and phosphatidylserine (POPS). Anionic phospholipids with small headgroups preferentially stabilize the complex **(C, D)** Sharpened cryo-EM maps for Erv14-Qdr2 reconstituted in lipid nanodiscs containing either one Erv14 molecule (ND1; contour level 0.34) or two Erv14 molecules (ND2; contour level 0.3) superimposed with density corresponding to the nanodisc (contour level ∼0.08, unsharpened maps). **(E)** Structural comparison of detergent and nanodisc complexes, revealing that Erv14-1 adopts a distinct position in the nanodisc structures, undergoing an approximately 27° rotation relative to Qdr2. The rotation axis is highlighted. Helices are labelled. **(F)** Surface representation of the ND2 complex showing an expanded lipid-filled luminal interface formed between Erv14 and Qdr2 following repositioning of Erv14-1. **(G)** Close-up of the luminal interface showing CHS-4-5 bridging the hydrophobic surfaces of Qdr2 and Erv14-1 following receptor rearrangement. Cryo-EM lipid density is contoured at 0.22 (sharpened map). **(H)** Co-immunoprecipitation analysis of Erv14 interface mutants. Mutation of the conserved hydrophobic anchor F18 markedly reduces association with Qdr2, whereas disruption of individual electrostatic interactions has little effect. Combined mutation of Site 1 3M (D29A, E31A, D33A) and Site 4 4M (H22A, D29A, E31A, D33A) also reduces association with Qdr2.

To explore how lipids influence the Erv14-Qdr2 interface, we determined cryo-EM structures of the complex reconstituted in lipid nanodiscs. Consistent with the biochemical data, the complex was less stable in nanodiscs, with partial dissociation of Erv14 during reconstitution (Fig. S4A). Cryo-EM analysis resolved two major populations containing either one Erv14 molecule bound to Qdr2 in a similar position to the LMNG structure (ND1; 2.85 Å), or two Erv14 molecules bound to Qdr2, with the second occupying a new site on the transporter (ND2; 2.76 Å) (Fig. 2C, D, Fig. S4 & Table S1). Superposition of the nanodisc and detergent structures showed that Qdr2 remains essentially unchanged (r.m.s.d. < 0.5 Å over 492 C_α_ atoms), whereas the receptor bound at the TM8, TM9 and TM12 interface (Erv14-1) adopts a distinct orientation, rotating by ∼27° relative to Qdr2 and reducing the buried interface to ∼400 Å² (Fig. 2E & Table S2).

The LPA-binding site is preserved in both nanodisc structures, with the lipid coordinating R365 and Q412 on Qdr2, together with Site 4 (K43) on Erv14-1 (Fig. S5A-C). CHS1 also occupies the same interfacial site around F18, although it is buried more deeply than in the detergent structure and forms an additional hydrogen bond with H22 in both nanodisc structures (Fig. S5D). By contrast, repositioning of Erv14-1 creates space on the luminal side of the interface, which is now occupied by two additional CHS molecules (CHS4-5). These bridge TM9 and TM10 of Qdr2 with TM1 and TM4 of Erv14 through hydrophobic interactions (Fig. 2F, G). Despite these changes, the D29-R525 salt bridge and the conserved F18 ‘knobs-in-holes’ interaction with TM9 are maintained (Fig. S5E, F), indicating that these interactions form the principal anchor points between receptor and cargo. Consistent with this model, mutation of F18 to arginine markedly reduced association with Qdr2 (Fig. 2H). In contrast, disruption of the electrostatic interface required combined mutation of Site 1 (D29A, E31A and D33A; 3M) together with H22 in Site 4 (4M), whereas individual mutations had little effect, indicating that electrostatic interactions are functionally redundant for cargo engagement.

### Structural basis for cornichon-driven membrane remodelling

An intriguing feature of the nanodisc structures was the presence of a second Erv14 receptor bound to Qdr2 (Erv14-2). Repositioning Erv14-1 creates space for this second cornichon to dock onto Qdr2 (Fig. S5G). As with Erv14-1, the second receptor engages the transporter through TM1 and TM4; however, it contacts only the extended TM8 helix of Qdr2 (Fig. 3A), burying ∼260 Å² of surface area, compared with ∼400 Å² in Erv14-1 (Table S2). Site 1 is the principal interaction site, with D29 and Y34 engaging R364 via salt-bridge and cation-π interactions, complemented by interactions between K368 and Y34 (Fig. 3B). The remaining interface is mediated by lipids that bridge Erv14-2 and Qdr2 (CHS6-7; ERG). The lysolipid acyl chain extends towards Erv14-2, where it contacts two CHS molecules (CHS6-7), one of which is also present in the ND1 structure (Fig. S5B, C). An ergosterol molecule packs adjacent to F18 (Fig. 3C), while the two CHS molecules adopt orientations tilted by ∼45° relative to the membrane normal (Fig. 3D). Notably, interactions involving F18 in Erv14-2 are mediated exclusively through lipids, highlighting the adaptability of the receptor-cargo interface.

**Fig. 3.**
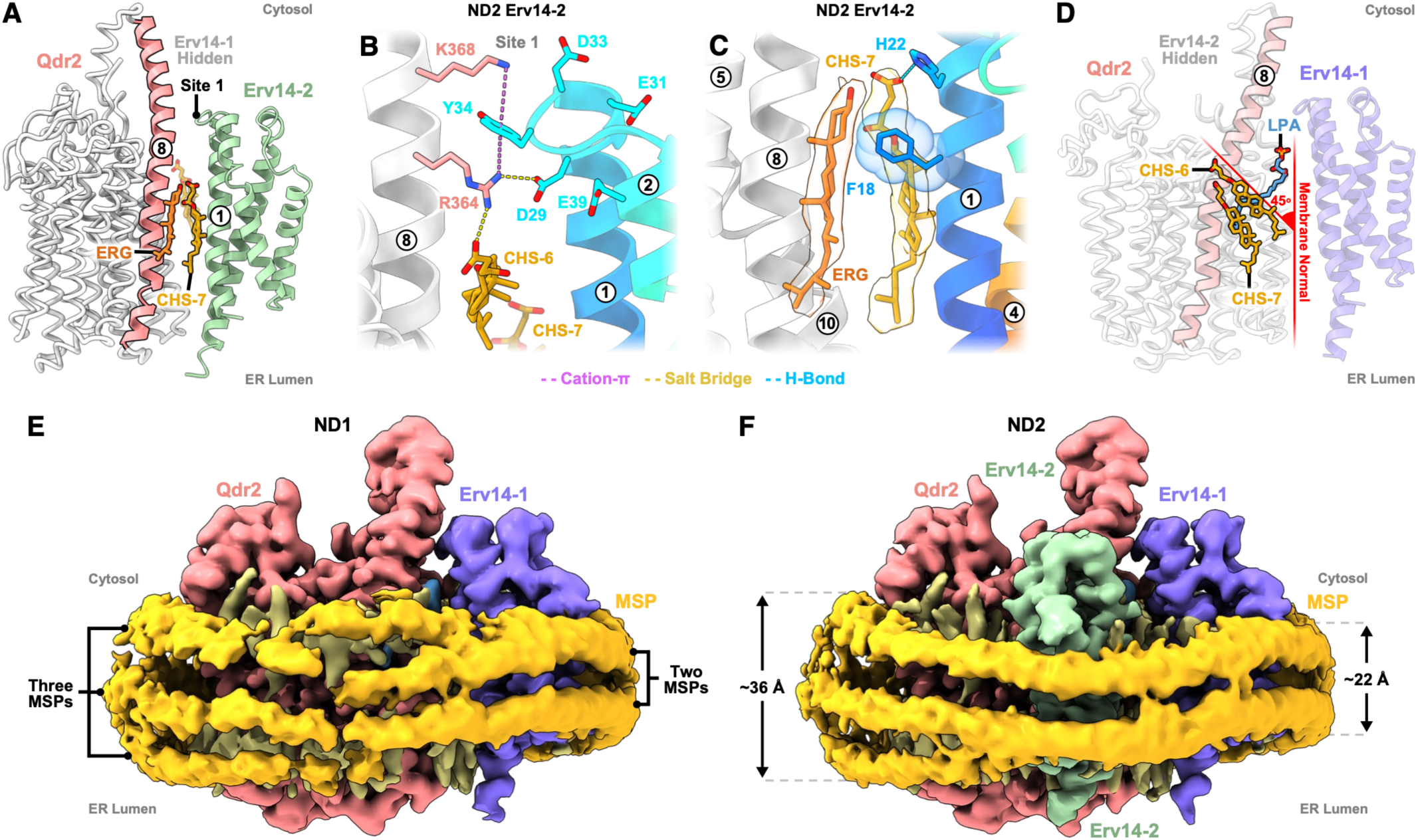
Recruitment of a second receptor buffers membrane thickness around Qdr2 to accommodate hydrophobic mismatch. **(A)** Overview of the Erv14-Qdr2 interface highlighting the extended Qdr2 TM8 and the extensive lipid-mediated interactions bridging the remaining receptor-cargo contacts. **(B)** Close-up of the Erv14-2 interaction site showing direct protein-protein contacts between Site 1 residues of Erv14 and Qdr2. **(C)** Close-up of Erv14-2 F18 packing against CHS7 and ERG at the Qdr2 interface. Cryo-EM lipid density is contoured at 0.23 (sharpened map). **(D)** Tilting of lipids (CHS6-7) relative to the membrane normal (red) at the Erv14-2-Qdr2 interface. These also pack against the acyl chain of LPA. Erv14-2 hidden. **(E, F)** Recruitment of a second Erv14 molecule into the ND2 complex reduces the membrane thickness from ∼36 Å to ∼22 Å and produces a more compact MSP belt around the transporter, consistent with Erv14 buffering hydrophobic mismatch within the membrane. Unsharpened cryo-EM maps shown, contoured at 0.11.

Comparison of detergent and nanodisc structures shows that receptor remodelling occurs primarily through rigid-body repositioning of the Erv14 receptors around Qdr2, rather than through conformational changes within the proteins (Fig. S5G & S6). Conserved lipid-binding sites are preserved, whereas protein contacts are redistributed to form smaller, more polar interfaces reinforced by hydrogen-bonding and electrostatic interactions (Table S2). These observations indicate that the Erv14-Qdr2 interfaces can remodel in response to changes in membrane architecture without large structural rearrangements.

How long transmembrane helices can be accommodated within the thin ER membrane remains an outstanding question in membrane protein biogenesis (*13, 15*). In ND1, three membrane scaffold protein (MSP) layers surround the hydrophobic surface of Qdr2, whereas only two are evident around Erv14 (Fig. 3E, F), indicating that recruitment of the second cornichon in ND2 reduces the effective membrane thickness surrounding the transporter. These structures provide direct experimental evidence that Erv14 buffers hydrophobic mismatch around Qdr2 and are consistent with the molecular dynamics simulations of Li *et al.* (*48*), which predict that Erv14 modulates local bilayer thickness. Our cryo-EM structures now provide direct structural evidence for this membrane buffering mechanism. The asymmetric architecture and hydrophobic surface of Erv14 likely underlie its ability to buffer membrane thickness around the receptor-cargo complex and demonstrate that multipass membrane proteins can recruit multiple cornichon receptors to facilitate their accommodation in different lipid environments within the cell.

### The second cornichon receptor promotes ER exit

To determine how the Erv14-Qdr2 interface contributes to ER export, we monitored the localisation of C-terminally GFP-tagged Qdr2 at the plasma membrane in yeast (*15, 49*). Mutation of the conserved hydrophobic anchor F18 to alanine, aspartate or arginine abolished Qdr2 trafficking to the plasma membrane (Fig. 4A-C & Fig. S7A), consistent with the reduced interaction observed in the pull-down assays (Fig. 2H). In contrast, mutation of individual electrostatic contacts at Site 1 (D29A) or Site 4 (H22A and K43A), or disruption of Site 1 alone (D29A/E31A/D33A; 3M), had little effect on trafficking. Only combined disruption of Site 1 and Site 4 (H22A/D29A/E31A/D33A; 4M) prevented Qdr2 export, reinforcing that electrostatic interactions are functionally redundant, whereas F18 forms the principal anchor for cargo recognition.

**Fig. 4.**
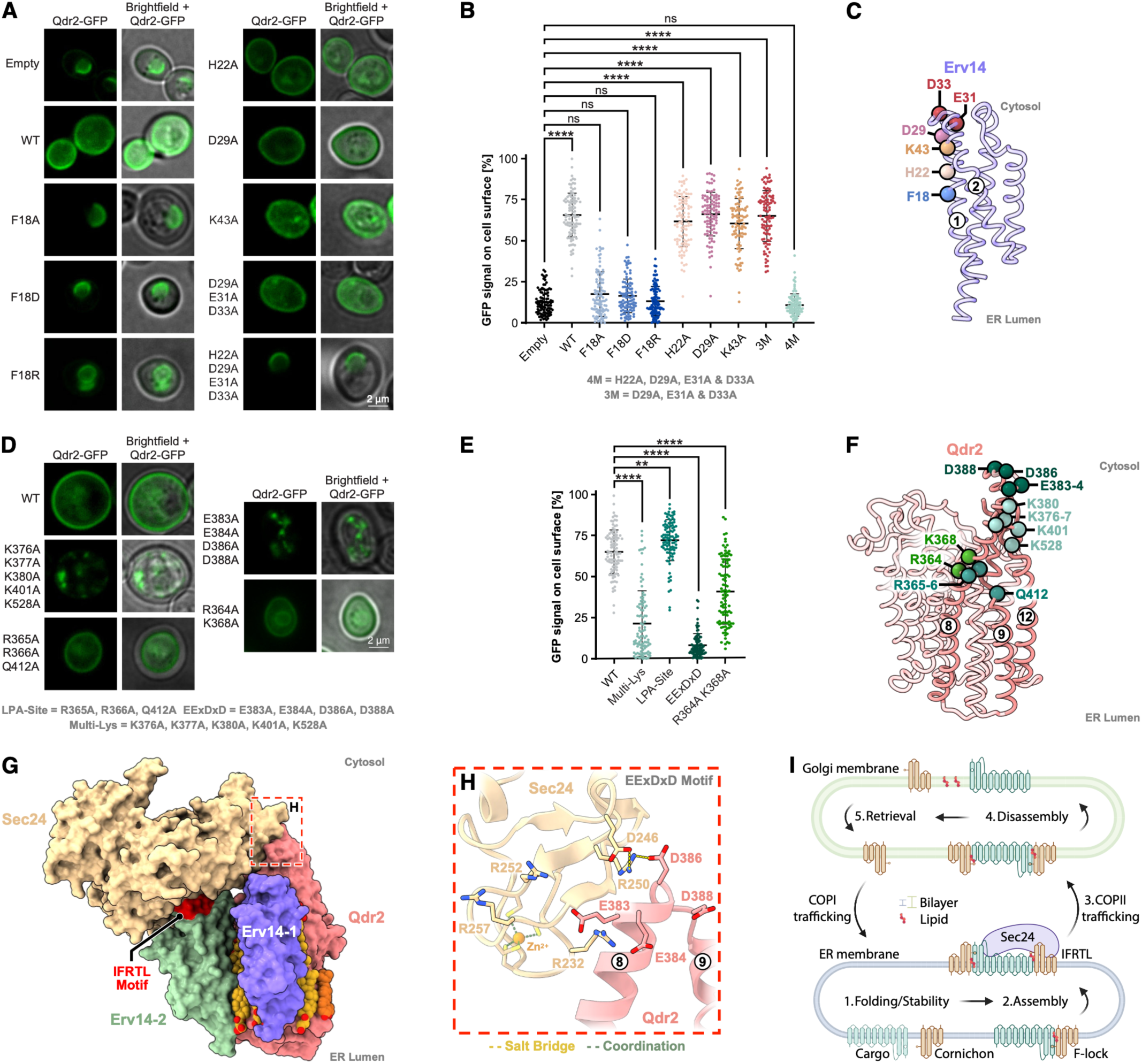
Distinct receptor interaction sites coordinate cargo maturation and Sec24 engagement. **(A, B)** Plasma membrane localisation of GFP-tagged Qdr2 expressed in a ΔErv14 yeast strain complemented with wild-type or mutant Erv14. Representative fluorescence micrographs are shown alone and merged with brightfield micrographs **(A)** and quantification of plasma membrane localisation **(B)**. **(C)** Structure of Erv14 indicating the location of the interface mutations. **(D, E)** Representative fluorescence micrographs are shown alone and merged with brightfield micrographs **(D)** and quantification of plasma membrane localisation **(E)** for Qdr2 mutants. **(F)** Structure of Qdr2 indicating the location of the interface mutations. **(G)** Model of the Erv14-Sec24 complex generated with AlphaFold3 and superimposed onto the ND2 structure, showing how Erv14-2 can recruit Sec24 without steric clash with Erv14-1 or Qdr2. The conserved IFRTL motif is positioned adjacent to the Sec24 D-site. **(H)** Close-up of the predicted interface between the conserved acidic motif within TM8-TM9 of Qdr2 and a basic surface on Sec24, identifying a candidate cargo-binding site involved in COPII recruitment. **(I)** Proposed model for cornichon-mediated ER export of multipass membrane proteins. (1) The first cornichon receptor functions as a membrane chaperone, stabilizing cargo folding within the thin ER membrane through conserved protein–protein and protein–lipid interactions with a key role for the TM1 phenylalanine (F-lock). (2) Following cargo maturation, recruitment of a second cornichon receptor buffers hydrophobic mismatch and promotes engagement of Sec24, driving assembly of the COPII coat. (3) The receptor-cargo complex is packaged into COPII vesicles and exported from the ER. Changes in membrane composition and thickness following ER exit promote dissociation of the cornichon receptors (4), which are subsequently retrieved to the ER by COPI-mediated transport (5).

We next turned our attention to the cargo, Qdr2, which exhibits unusual structural features for an MFS transporter. Intriguingly, the LPA lipid and Erv14-1 interaction sites sit near an unwound region of TM9, which is itself an unusually long and highly charged TM helix (Fig. 1D). This raised the question of whether Erv14 may function as a membrane chaperone for polytopic membrane proteins. Monitoring membrane protein expression levels using a C-terminal GFP-tagged protein is a well-established method for studying membrane protein biogenesis (*49, 50*). Consistent with Erv14 having a role in protein folding (*51*), expression of Qdr2-GFP increased in the presence of wild-type Erv14 but was reduced by binding-deficient mutants, F18R and 4M (Fig. S7B, C). Mutation of the positively charged surface on TM8-TM9 (K376A/K377A/K380A/K401A/K528A; Multi-Lys) also impaired protein stability and trafficking (Fig. 4D-F & Fig. S7D-F & K), supporting a role for this interface in cargo maturation. Mutation of the LPA site (R365A/R366A/Q412A; LPA-Site) resulted in correctly localised Qdr2 (Fig. 4D-E); however, expression was much slower than that of WT Qdr2 (Fig. S7B, C), indicating that LPA binding promotes stabilisation of Qdr2 during biogenesis. Loss of both the LPA binding site and Erv14 results in a severely compromised Qdr2 protein (Fig. S7I-J), indicating that both factors are important for efficient Qdr2 synthesis and folding.

To investigate how cornichon receptors recruit the COPII coat, we modelled the Erv14-Sec24 complex using AlphaFold3 (AF3) (49). Although a 2:1:1 Erv14-Qdr2-Sec24 model failed to recapitulate the experimentally observed assembly, a 1:1:1 model positioned Erv14 at the second receptor-binding site identified in the ND2 structure (Erv14-2). Superposition of the predicted Erv14-Sec24 complex onto the ND2 structure showed that Sec24 can be accommodated without steric clash with Erv14-1 or Qdr2 (Fig. 4G & Fig. S8A). Consistent with this model, mutation of the Erv14-2 interaction site on Qdr2 (R364A/K368A) (Fig. 3B) impaired trafficking without affecting protein stability (Fig. 4D, F & Fig. S7G), indicating that recruitment of the second Erv14 receptor promotes ER export by recruiting Sec24.

Several cargo-binding sites have been identified on Sec24, many of which are conserved among human paralogues (*10*). These sites, known as the A-, B-, C-, and D-sites, recognise distinct ER export motifs and function independently (*9, 52, 53*). The predicted AF3 complex places the conserved I^97^FRTL export motif of Erv14-2 adjacent to the Sec24 D-site, the cargo-recognition pocket that binds IFXL export motifs (*10*). Specifically, F576 from Sec24 packs against the cavity formed near the I^97^FRTL loop on the cornichon receptor, and R578 from the D-site lies adjacent to Site 1 in Erv14-2. Additionally, R99 from the IFR^99^TL motif forms a salt bridge to E661 on Sec24 (Fig. 4G & Fig. S8B). The AF3 model also positions an acidic motif (E^383^ExDxD) within TM8-TM9 of Qdr2 near a conserved basic surface on Sec24 that contains R232, R250, R252, and R257 (Fig. 4H). This cluster of basic side chains on Sec24 has not previously been identified as a potential cargo-binding motif (*10*) but is conserved in human Sec24 paralogues (Fig. S8C). Mutation of the E^383^ExDxD motif to alanine abolished Qdr2 trafficking while preserving interaction with Erv14 (Fig. 4D-F & Fig. S7H, K), supporting a role for this previously unrecognized Sec24 surface in cargo engagement during COPII vesicle assembly.

Our data, together with a complementary model proposed by Li *et al*. (*48*), suggest that cargo engagement drives the formation of an export-competent Erv14 assembly (Fig. 4I). Rather than recognising cargo through a single invariant interface, Erv14 engages Qdr2 through reconfigurable interaction networks, while preserving structural lipid interactions. This engagement is driven in part by the conserved phenylalanine on TM1 (F-lock). We propose that cornichon receptors perform sequential functions during membrane protein biogenesis: one receptor stabilises cargo during folding, whereas recruitment of additional receptors promotes membrane adaptation and COPII assembly, thereby coupling protein quality control to selective ER export. This model provides a structural framework for understanding how cornichon receptors coordinate membrane protein biogenesis with selective cargo trafficking across eukaryotes.

## Data and materials availability

Atomic coordinates for the Erv14-Qdr2 structures have been deposited in the Protein Data Bank under accession codes 32PZ (LMNG: CHS structure), 32QB (ND1), and 32QA (ND2).

The cryo-EM maps have been deposited in the Electron Microscopy Data Bank (EMDB) under accession codes:

[https://www.ebi.ac.uk/pdbe/entry/emdb/EMD-59064] (PDB:32PZ).

[https://www.ebi.ac.uk/pdbe/entry/emdb/EMD-59066] (PDB:32QB).

[https://www.ebi.ac.uk/pdbe/entry/emdb/EMD-59065] (PDB:32QA).

## Acknowledgements

This research was supported by Wellcome awards (219531/Z/19Z & 320911/Z/24/Z) to SN, and a BBSRC award (BB/Z517215/1) to SN and JLP and by the Intramural Research Program of the NIH, National Institute of Neurological Disorders and Stroke (ZIA NS003139) to LRF. The contributions of the NIH authors were made as part of their official duties as NIH federal employees, are in compliance with agency policy requirements, and are considered Works of the United States Government. However, the findings and conclusions presented in this paper are those of the authors and do not necessarily reflect the views of the NIH or the U.S. Department of Health and Human Services. This work utilized the computational resources of the NIH HPC Biowulf cluster. Fig. 4I was created in https://BioRender.com. The authors wish to thank Carolina Yanez-Dominguez, Universidad Nacional Autónoma de México, for initial discussions at the start of this project. Dušan Živković and Jani R. Bolla, University of Oxford, for assistance with native mass spectrometry and lipidomics and Maya Schuldiner, Weizmann Institute of Science, for sharing yeast strains. Cryo-EM data were collected at the Central Oxford Structural Molecular Imaging Centre (COSMIC, University of Oxford) and fluorescence microscopy undertaken at the Micron Bioimaging Facility (University of Oxford). We gratefully acknowledge staff from both facilities for their support and assistance in this work.

## Author contributions

JLP & SN conceived the project. JT, OA, SH, GK, and JLP performed all cloning, protein preparation and biochemical experiments. OA and JT performed all cryo-EM sample processing, data collection and image analysis. JT, OA, JLP and SN constructed the atomic models. MB and SH performed all fluorescent microscopy data acquisition and analysis. JT, NB and LRF performed all molecular dynamics simulations and analysis. OP provided yeast strains. OA, JLP and SN analysed the data and wrote the paper.

