## Supplementary Material for "Cornichon receptors couple membrane adaptation to cargo selection during ER export"

**Supplementary Information**

**Materials & Methods**

**Figures S1-S8**

**Table S1-S2**

### Materials and Methods

#### Erv14-Qdr2 complex expression and purification

The gene encoding full-length *Saccharomyces cerevisiae* (Sc) Qdr2 (residues 1-542, Uniprot: P40474) was synthesised as a DNA fragment and cloned into the pDDGFP-Leu2D vector (Addgene 102334) (*1, 2*). Erv14 was amplified from Sc genomic DNA with a C-terminal FLAG tag added via PCR and cloned into YEplac112 (BamHI/PstI), which contained a GAL promoter (SacI/BamHI) and a GBT9 terminator (PstI/SphI). Both plasmids were transformed into the yeast strain BJ5460, and the strain was cultivated in synthetic complete medium minus leucine and tryptophan, supplemented with 2% (w/v) D-glucose. An overnight culture was diluted tenfold into medium containing 2% (v/v) lactate, and expression was induced after 14 h by adding 2% (w/v) galactose. The yeast cells were harvested after a further 22 h, and the membranes were prepared. Membrane preparation consisted of lysing the cells under high pressure (38 kpsi using a Constant Systems continuous-flow cell disruptor). Unbroken cells and cellular debris were pelleted at 30,000 g for 25 min at 4 °C. Membranes were harvested by centrifugation at 235,000 g for 1.5 h at 4 °C and washed once with 20 mM HEPES pH 7.5, 1 M potassium acetate. The membranes were resuspended in PBS and snap-frozen at – 80 °C for storage until required.

For purification, thawed membranes (~12 g wet weight) were solubilised in 130 ml of buffer containing 1 x PBS supplemented with 150 mM NaCl, 10% glycerol, 1% (w/v) Lauryl Maltoside Neopentyl Glycol (LMNG) detergent (Anatrace Cat. No. NG318), and 0.1% (w/v) Cholesteryl hemisuccinate (CHS) (Merck Cat. No. C6512) for 1.5 h under gentle agitation on a magnetic stir plate. All purification steps were carried out at 4 °C. Following solubilisation, the insoluble material was removed by centrifugation at 235,000 g for 1 h. 25 mM imidazole was added to the supernatant, which was then batch-bound to HisPur-Ni-NTA resin (Fisher Scientific Cat. No. 10038124) for 4 h with gentle stirring. The resin was loaded onto an Econo-column (BioRad) and washed with eight column volumes (CVs) of buffer (1X PBS, 10% glycerol, 150 mM NaCl with 0.1% LMNG: CHS (10:1)) containing 25 mM imidazole, followed by 12 CVs of wash buffer containing 40 mM imidazole. Protein was eluted with four CVs of buffer containing 250 mM imidazole and dialyzed overnight with TEV protease. Following TEV cleavage, 10 mM imidazole was added to the dialysate, and the mixture was then passed through a 5 ml HisTrap column (Cytiva) in a reverse IMAC purification step. The flow-through was incubated in batch with Pierce Anti-DYKDDDDK affinity resin (Thermo Fisher Scientific) for 2 h and then subjected to gravity flow. The flow-through was collected and incubated with additional Pierce Anti-DYKDDDDK affinity resin (Thermo Fisher

Scientific) for a further 1.5 h to ensure capture of all protein. The resins were washed with 30 CV of buffer (containing 0.0015% LMNG: CHS) and then eluted with 3x FLAG peptide for 1 h. The two elution pools were combined, concentrated to 500 µL using an Amicon Ultra-2 Centrifugal Filter (30 kDa MWCO, Millipore) at 4 °C, and subjected to size exclusion chromatography (Superdex 200 Increase 10/300 GL column) in a buffer consisting of 20 mM Tris pH 7.5, 150 mM NaCl, 0.0015% LMNG: CHS. The peak fractions containing Qdr2 and Erv14 were pooled, concentrated to 6.1 mg/ml and used to prepare cryoEM grids. For nanodisc preparation, the protein was purified as above, using DDM: CHS (5:1), 1% for solubilisation, 0.1% for washing, and 0.03% in the final SEC chromatography.

**Nanodisc reconstitution**

The Erv14-Qdr2 complex purified in DDM:CHS was reconstituted into MSP1E3D1 lipid nanodiscs using asolectin supplemented with 7% (wt/wt) POPS via the Bio-Bead method (*3*). In brief, nanodiscs were assembled at a 1:5:100 protein: lipid: MSP ratio in a final volume of 3.7 ml of reconstitution buffer (20 mM HEPES pH 7.5, 150 mM NaCl, 15 mM sodium cholate) and at a final Erv14-Qdr2 complex concentration of 2 µM. Lipid was pre-solubilised in 3x molar excess sodium cholate. After 1.25 h incubation on ice, detergent removal was achieved in two consecutive Bio-Bead (SM-2 Resin, Bio-Rad) incubation steps, the first for 1.5 h and the second overnight, and both at 4 °C with gentle rotation. Each round used a total of ~740 µl of Bio-Beads, split among multiple 1.5 ml Eppendorf tubes. Bio-Beads were extracted, and the assemblies then ultracentrifuged (100,000 g, 30 min, 4 °C). The recovered supernatant was pooled, concentrated to 500 µl (Vivaspin 50 kDa MWCO, Sartorius) and subjected to size exclusion chromatography (Superdex 200 Increase 10/300 GL column (Cytiva)) in 20 mM HEPES pH 7.5, 150 mM NaCl buffer. Fractions containing the complex and MSPs were concentrated (Vivaspin 50 kDa MWCO, Sartorius) to 3.97 mg/ml and immediately used for cryo-EM grid preparation.

**Cryo-EM sample preparation and data acquisition**

3 µl of protein (LMNG: CHS samples at 2, 4, or 6.1 mg/ml; nanodiscs at 3.97 mg/ml) was applied to freshly plasma-cleaned holey carbon grids (Quantifoil R1.2/1.3, 300-mesh Cu or Au). After 7 s of adsorption (LMNG: CHS samples) or immediately for the nanodiscs, grids were blotted (2.5-6 s, blot force +5/7, 100% humidity, 4 °C) and plunge-frozen in liquid ethane using a Vitrobot Mark IV (Thermo Fisher Scientific). Data were collected in counted super-resolution mode (bin2) on a Titan Krios G3 (Thermo Fisher Scientific), equipped with a

BioQuantum imaging filter (Gatan) and a K3 direct-detection camera (Gatan), operating at 300 kV and 105,000x magnification (pixel size 0.832 Å). From two collections each, a total of 58,208 and 49,836 movies were acquired (defocus range -2.2 to -1.0 µm) for the LMNG: CHS and nanodisc samples, respectively. Across all collections, the total dose spanned 39.1-41.1 $e^{-}/Å^{2}$ and the exposure time 1.8-2.4 s. Representative micrographs are shown in Fig. S1B and Fig. S3B.

**Cryo-EM data processing**

Schematics summarising the data processing workflows are provided in Fig. S1C (LMNG: CHS sample) and Fig. S4C (nanodisc sample). For all datasets, patched motion correction, patched contrast transfer function (CTF) estimation, particle picking, and particle extraction (300-px box) were performed on the fly using SIMPLE 3.0 (*4*). Unless stated otherwise, all subsequent processing was performed in cryoSPARC (*5*). Throughout processing, all refinement reference maps were lowpass filtered to at least 8 Å, and global resolutions estimated via gold-standard Fourier shell correlations (FSCs) using the 0.143 criterion. For the LMNG: CHS sample, pre-processing of the first collection (26,205 micrographs) yielded an initial stack of 9,825,631 particles. Following several 2D classification rounds, discarding poor-quality classes after each, the remaining 2,358,574 particles underwent multi-class (K=4) *ab initio* reconstruction. This generated a single class (627,225 particles) displaying clear features of the Erv14-Qdr2 complex incorporated into a detergent micelle. Further sorting of these particles via multiple rounds of heterogeneous refinement gave a stack (358,056 particles) producing a 4.07 Å map on non-uniform (NU) refinement. These particles were carried forward for Bayesian polishing in RELION 3.1.3 (*6, 7*), using the csparc2star script to move between programmes (*8*). After additional cleaning by 2D classification, heterogeneous refinement, and masked 3D classification (K=2, filter resolution ~3.2 Å, Erv14-Qdr2 protein mask), the retained 245,348 particles returned a 3.43 Å reconstruction on NU refinement. Using this as the target volume, heterogeneous refinement was then employed to recover Erv14-Qdr2 particles from both collections (58,208 micrographs total; input stack 10,276,294 particles, post-2D classification). Subjecting the recovered 3,276,313 particles to two rounds of masked 3D classification (K=10/2, filter resolution 2.75/2.5 Å) yielded 647,621 particles NU refining to 2.96 Å. These were then Bayesian polished as above. Following final rounds of 2D classification and heterogeneous refinement, interspersed with global and local CTF refinement, a final stack of 376,696 particles yielded a 2.83 Å reconstruction on successive NU and masked local refinement.

For the nanodisc sample, pre-processing of the initial collection (15,426 micrographs) produced a stack of 10,570,860 particles. After multiple rounds of 2D classification, the retained 2,760,768 particles were subject to multi-class (K=5) *ab initio* model generation. A single class (838,309 particles) clearly corresponded to nanodisc-reconstituted Erv14-Qdr2, and further processing gave a 3.34 Å volume following NU refinement of a 380,248-particle subset. Erv14-1-Qdr2 was well resolved in this map, with partial density for Erv14-2 already apparent. This reconstruction was next used as the target volume in a heterogeneous refinement aimed at recovering Erv14-Qdr2 particles from the entire dataset (49,836 micrographs total; input stack 15,266,009 particles, post-2D classification). The surviving 5,049,982 particles were further cleaned by 2D classification followed by iterative rounds of heterogeneous refinement, eventually leaving a stack of 1,262,651 particles. NU refinement of these yielded a 2.89 Å volume with improved, but still partial, density for Erv14-2. Next, in an attempt to separate complexes with differing Erv14 occupancy, focused 3D classification (K=4, filter resolution 10 Å) was trialled with Erv14-2-only, Erv14-2-Qdr2, and Erv14-1-2-Qdr2 protein masks. All three classifications identified volumes containing either two well-resolved Erv14 molecules, at the Erv14-1 and Erv14-2 sites, or only a single copy bound at the Erv14-1 site. The best singly and doubly Erv14-bound classes from each classification were then pooled, with duplicate particles removed, for subsequent independent processing. This gave 534,562 and 468,278 particles for the double-Erv14 (ND2) and single-Erv14 (ND1) states, respectively. Each stack was next subject to a single round of 3D classification (K=3, filter resolution ~2.6 Å, entire complex masks) to identify particle subsets yielding improved resolution. These were then advanced for Bayesian polishing in RELION 5.0 (*6, 7*) and, after a last round of 2D classification, gave final reconstructions of 2.76 Å (ND2, 207,007 particles) and 2.85 Å (ND1, 168,498 particles) following NU and local refinement.

**Model building and refinement**

The AlphaFold (*9*) models of Erv14 (ID AF-P53173-F1) and Qdr2 (AF-P40474-F1) were manually fitted in the maps in ChimeraX v. 1.10.1 (*10*) to generate the initial models. After fitting, the models were manually readjusted using COOT (*11*) and refined using PHENIX(*12*) and ISOLDE (*13*). The figures depicting the molecular structures were prepared using ChimeraX v. 1.10.1.

**Coarse-grained representation molecular dynamics simulations**

Molecular dynamics simulations were performed to examine lipid-protein interactions in the cryo-EM model of the detergent-solubilised Erv14-Qdr2 complex. The atomic structure was coarse-grained using the martinize tool (*14*), embedded in a 2:1 POPE: POPG bilayer, and solvated, resulting in a box of 200 × 200 × 200 Å. The system was charge-neutralised by adding $Na^+$ and $Cl^-$ ions to a concentration of 150 mM. In preparation for the production steps, a 15,000-step energy minimisation using the steepest-descent method was performed, followed by a 5 ns equilibration to reach the desired temperature and pressure. All simulations were performed with GROMACS v 2024.1 (*15, 16*) using the MARTINI 2.2 forcefield (*14*). Temperature and pressure were maintained at 303.15 K and 1 bar, respectively, using the velocity-rescale thermostat and the Parrinello-Rahman semi-isotropic barostat. Inter- and intra-protomer protein structures were retained using constraints of 10 kJ/mol–1 between all backbone beads within 20 Å. Production runs were 20 µs long.

**Analysis of molecular dynamics simulations**

Coarse-grained trajectories were prepared for spatial analysis by centring the protein, placing all molecules within the box, and then performing 2-dimensional fitting around the protein alpha-carbons. All analysis was then performed using the MOSAICS toolset (*17*). The mean lipid dwell times were measured using the 2d Kinetics tool, which uses Voronoi tessellations of the lipid molecules for each trajectory frame, defined around the GL1 and GL2 coarse-grained beads. The analysis monitors the Voronoi cells and records the dwell time for each lattice point as new cells are encountered. Similarly, the lipid density maps were generated using Lipid Density 3D with a 2.6 Å stamping radius around each coarse-grained bead. A value of 1 is stamped around each bead, and the lattice is normalised by the number of trajectory frames. The spatially resolved time-averaged atomic coordinates for the lipids were measured and plotted alongside the corresponding averages for the protein. This analysis used the Mean Lipid Coords tool.

**Co-immunoprecipitation assays**

Cultures were grown as detailed for the large-scale purification protocol in a final volume of 200 ml. After induction with galactose, the $OD_{600}$ of the culture was measured, and the equivalent of 180 ml of a culture at OD 3 was harvested. Yeast cells were lysed in PBS by vortexing with silica beads, and unlysed yeast were removed by centrifugation at 4,000 g for 30 seconds. Membranes were then harvested by centrifugation at >30,000 g for 1 h, resuspended in PBS, aliquoted into 3 aliquots, and snap-frozen in liquid nitrogen before storage at -80 °C. FLAG pulldowns were performed using one membrane aliquot (per condition), solubilised in 1% detergent in PBS to a final volume of 1 ml. The clarified material (30 min at 200,000 g) was incubated with FLAG resin (20 µl) for 2.5 hours on a rotating wheel at 4 °C. The resin was recovered by centrifugation (800 g for 30 s) and washed three times with 200 µl wash buffer (typically 0.1% detergent in PBS). Bound proteins were eluted by adding 25 µl of LDS buffer for 5 min, then subjected to Western blot analysis using an anti-FLAG antibody (Merck F1804 at 1:8000 dilution) for Erv14 detection and an anti-GFP antibody (Merck G1544 at 1:5000 dilution) for Qdr2 detection.

**Size exclusion chromatography analysis**

Yeast cultures containing WT Qdr2-GFP and WT Erv14-FLAG plasmids were grown as for the large-scale purification protocol, and membranes were prepared in the same way. For each lipid condition tested, a 4% LMNG stock made in 0.2 M Tris pH 8.1 was used to solubilise the lipid of choice at a ratio of 10:1 (detergent to lipid). 0.4 g of membrane was solubilised in PBS (total volume 5 ml) with a final detergent/lipid concentration of 1% for 1.25 h at 4 °C with rotation, and insoluble material was removed by centrifugation at 280,000 g for 25 minutes. FLAG resin (65 µl) was added for 2.25 h, then washed 4 times (2 x 0.2 ml and 2 x 0.25 ml) with PBS + 0.1% detergent. Bound protein was eluted with 0.4 ml of 3x FLAG peptide (0.8 mg/ml in PBS + 0.005% detergent) for 1 hour with rotation, and the resin was washed with 100 µl of the same buffer. Eluted material (0.5 ml) was subjected to SEC (Superdex 200) in buffer containing PBS + 0.003% detergent. The data were normalised to the Erv14-FLAG peak (elution volume of ~13.2 ml), which was set to 100, and the data shown in Fig. 2B relate to the complex peak height (~11.5 ml). Each detergent: lipid small-scale purification was repeated in triplicate, and representative traces are shown in Fig. S2E-G.

**Trafficking assays**

The gene encoding Erv14, with a C-terminal FLAG tag and including the genomic sequence 265 base pairs upstream and 242 base pairs downstream, was synthesised with SalI and BamHI restriction sites. This was cloned into the YCplac111 vector (minus leucine selection). Mutant variants were made by site-directed mutagenesis and verified by sequencing. The protein levels of each mutant were analysed by immunoblotting with an anti-FLAG antibody; 5 ml of the cultures used for the imaging (at an $OD_{600}$ of 0.8) were used to prepare membranes. The membranes were subjected to SDS-PAGE and Western blot analysis using anti-FLAG antibody (Merck F1804 at 1:6000 dilution). To allow visualisation of Qdr2, the WT QDR2 gene was cloned into the pDDGFP vector, which encodes a C-terminal GFP tag (minus uracil selection). This vector drives expression of the protein of interest under the galactose promoter, which was used to increase the amount of Qdr2 in the cell, allowing for quantification of its localisation (plasma membrane vs intracellular). Mutant variants of each gene were made by site-directed mutagenesis and verified by sequencing. *S. cerevisiae* yeast strain BY4741ΔErv14 (Horizon Discovery No. YSC6273-201936099) was transformed with combinations of the plasmids as required and selected on SC minus Leu/Ura agar plates. Cultures were grown overnight in selective medium supplemented with 2% glucose before being transferred to medium supplemented with 2% lactate. Qdr2 protein expression was induced with 2% galactose for 3 h. Cells were washed with PBS, resuspended in PBS, and placed on a freshly prepared agarose pad (20 mg/ml agarose in PBS) on a glass slide, then covered with a glass coverslip. Live cells were imaged with a 100X HC PL FLUOTAR Oil NA 1.3 (OM44) objective (Leica Microsystems, Germany) on a Leica Thunder inverted microscope (Leica Microsystems) with an sCMOS Hamamatsu Orca Fusion Digital C14440-20UP camera (Hamamatsu Photonics K.K., Japan) and LAS 3.10.1 software (Leica Microsystems). Cells were illuminated with a CoolLED pE-800 LED Light Engine (CoolLED, UK). Plasma membrane localisation was quantified using Fiji software (*18*). The total fluorescence of each cell was calculated within an ovoid shape (drawn in Fiji; yeast cells with altered morphology were excluded from the analysis) and compared with the total fluorescence within the cell. The ratio of intracellular to total fluorescence was calculated as a percentage, and this value was subtracted from 100 to determine the plasma membrane amount. The percentage of GFP fluorescence from the plasma membrane for each cell was plotted in GraphPad Prism v11 software (www.graphpad.com). To prepare the figures, Thunder computational clearing deconvolution algorithms were applied. Merged images of fluorescence micrographs and brightfield micrographs were prepared and cropped in LAS 3.10.1 to show individual cells, then pasted into Affinity Designer.

**Qdr2-GFP expression analysis**

The yeast strain and plasmids used for imaging analysis were used to quantify Qdr2 expression, measured by fluorescence from the C-terminal GFP tag. Yeast cultures were grown overnight in glucose, then back-diluted to an $OD_{600}$ of 0.25 in lactate and incubated for 14 h at 28 °C. After this time, the temperature was raised to 30 °C, and galactose was added to a final concentration of 1.5%. Both $OD_{600}$ and GFP fluorescence were monitored. Typically, at the time of induction, the cultures were at an OD of 1-1.4. For each time point, the equivalent of 2 $\times 10^8$ cells were harvested (in duplicate), resuspended in 100 μl of PBS, and fluorescence (excitation 488 nm, emission 510 nm, on a SpectraMax M3 fluorimeter – Molecular Devices) was measured, and an average was taken. To account for day-to-day variation, each mutant was repeated in triplicate on at least two separate days, and for each set of experiments, the positive control (WT Qdr2 with WT Erv14) and negative control (WT Qdr2 with the empty YCplac111 plasmid) were carried out in parallel. The data were plotted as percentages, using the highest expression value for WT Qdr2 with WT Erv14 in each experiment as 100%.

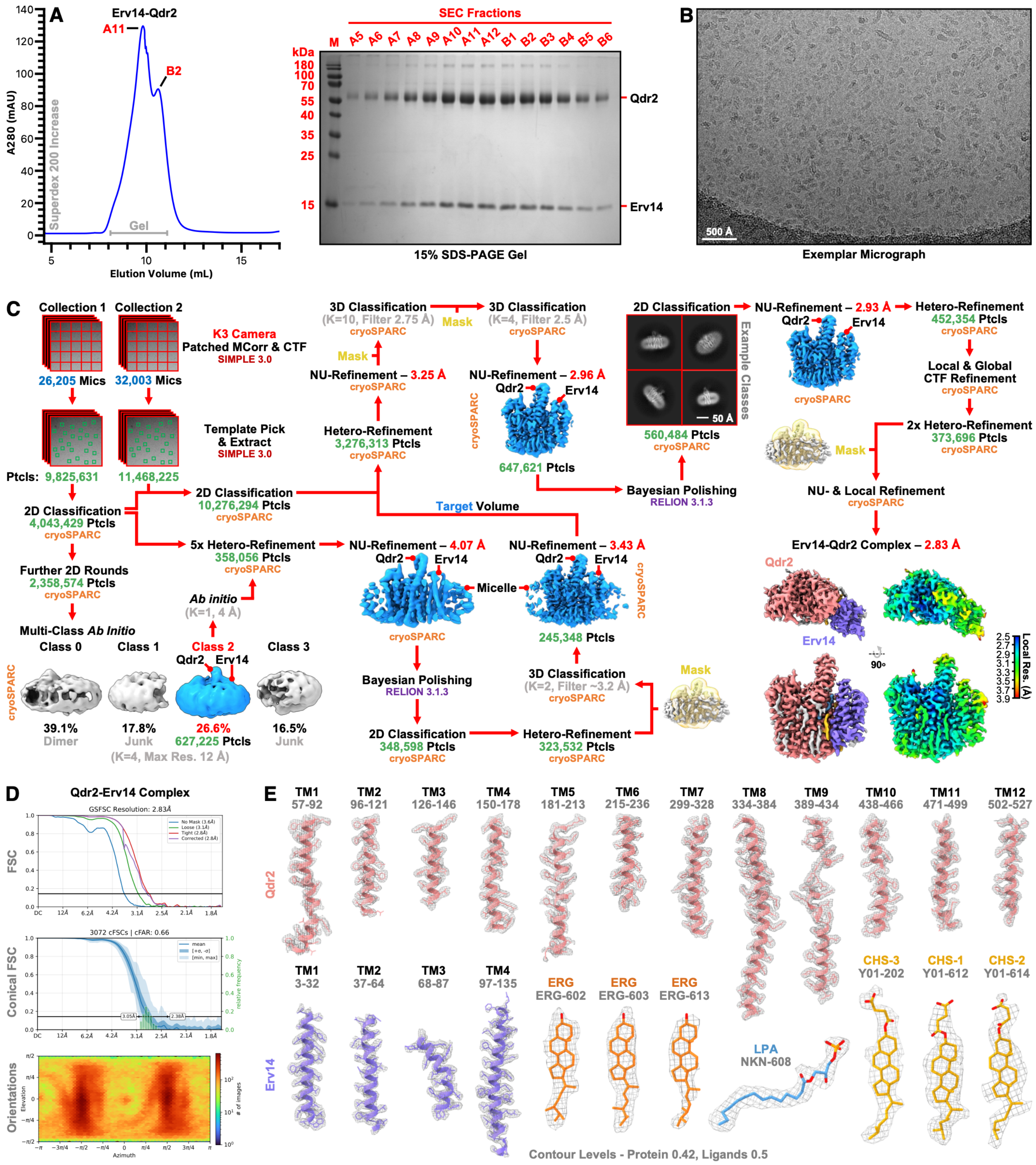


**Figure S1. Cryo-EM workflow, map resolution and model density fit for the Erv14-Qdr2 complex in LMNG: CHS. (A)** SEC profile of the purified Erv14-Qdr2 complex in LMNG: CHS; the grey bar indicates the fractions shown in the Coomassie-stained SDS-PAGE gel. (**B**) Representative micrograph. (**C**) Overview of the data acquisition and processing workflow, with the software packages used at each stage indicated. Local resolution estimates for the sharpened map (contour level 0.6) are also shown. (**D)** Gold-standard Fourier shell correlation (FSC) curves estimating global resolution (0.143 cut-off), alongside conical FSC and orientation plots for the final reconstruction. **(E)** Density fits for all transmembrane helices of both Qdr2 and Erv14, as well as all modelled lipids.

A

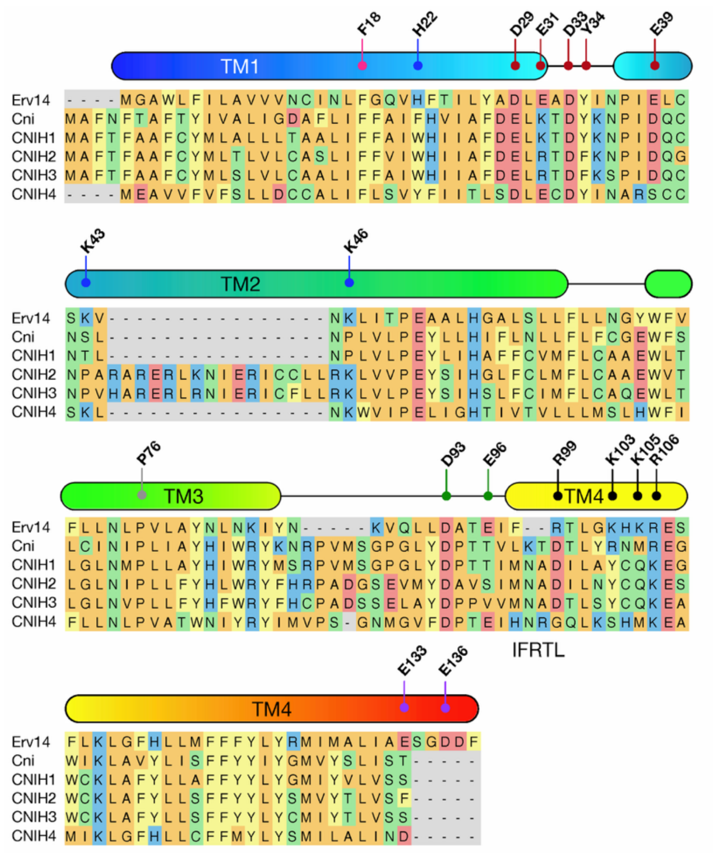


B

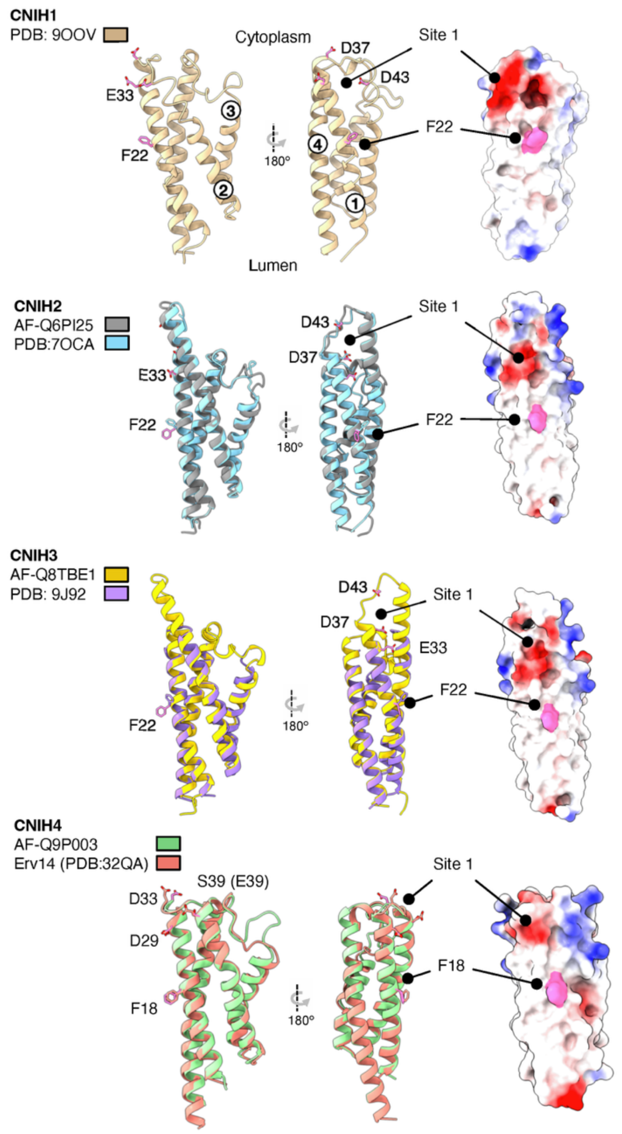


C

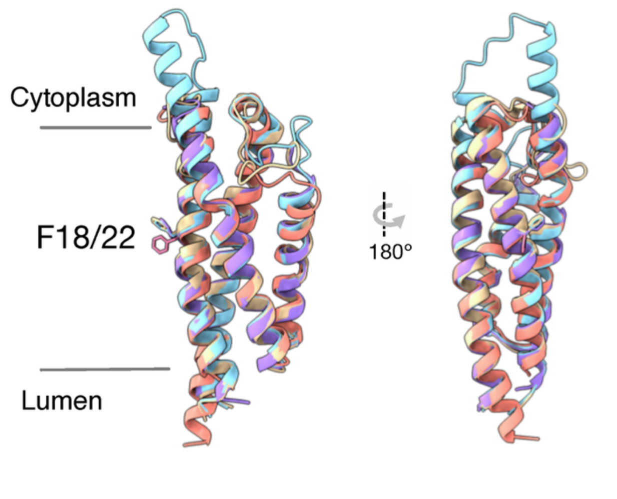


D

| | Erv14 (32QB) | CNIH1 (9OVV) | CNIH2 (7OCA) | CNIH3 (9J92) |
|---|---|---|---|---|
| Erv14 (32QB) | - | | | |
| CNIH1 (9OVV) | 1.24 | - | | |
| CNIH2 (7OCA) | 0.916 | 0.802 | - | |
| CNIH3 (9J92) | 0.946 | 0.832 | 0.642 | - |

E

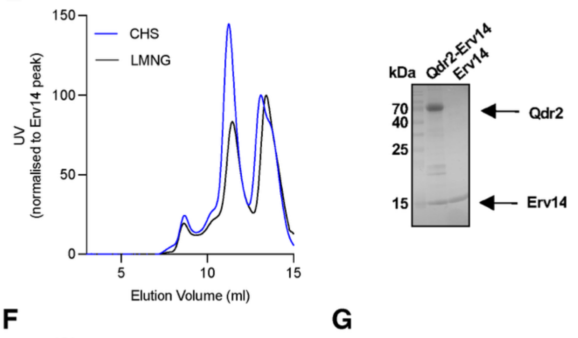


F

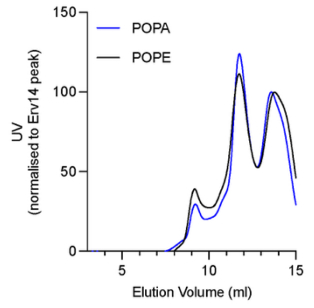


G

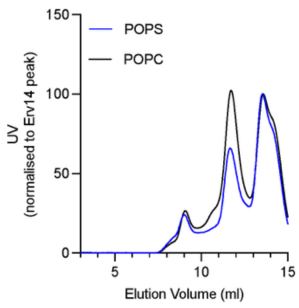

**Figure S2. Structural conservation supports a common mechanism of cornichon function. (A)** Structure-guided sequence alignment of *Saccharomyces cerevisiae* Erv14 (Uniprot: P53173) with the founding cornichon homologue Cni from *Drosophila melanogasta* (P49858) and the human CNIH1-4 homologues (O95406, Q6PI25, Q8TBE1, Q9P003). Conserved residues involved in cargo recognition, including F18 and charged interaction sites, are indicated. The IFRTL Sec24 export motif is highlighted. **(B)** Comparison of experimentally determined structures and AlphaFold models of mammalian CNIH1-3 with the Erv14 structure (PDB: 32QA). AlphaFold models were used where the PDB model was incomplete from the experimental structures. Cartoon and electrostatic surface representations show conservation of the overall four-transmembrane-helix architecture, asymmetric molecular shape, and the negatively charged Site 1 despite sequence divergence. **(C)** Structural superposition of Erv14 with mammalian CNIH proteins demonstrating conservation of the transmembrane architecture and the position of the conserved aromatic residue (F18/F22) implicated in cargo recognition. **(D)** Pairwise structural comparisons ($C_\alpha$ r.m.s.d.) between Erv14 and mammalian CNIH proteins demonstrate conservation of the cornichon fold. **(E)** Size-exclusion chromatography and SDS–PAGE analysis showing stabilization of the Erv14-Qdr2 complex by CHS. **(F)** Size-exclusion chromatography analysis showing stabilization of the Erv14-Qdr2 complex in the presence of POPA and POPE. **(G)** Size-exclusion chromatography analysis showing stabilization of the Erv14–Qdr2 complex in the presence of POPC and destabilisation in the presence of POPS.

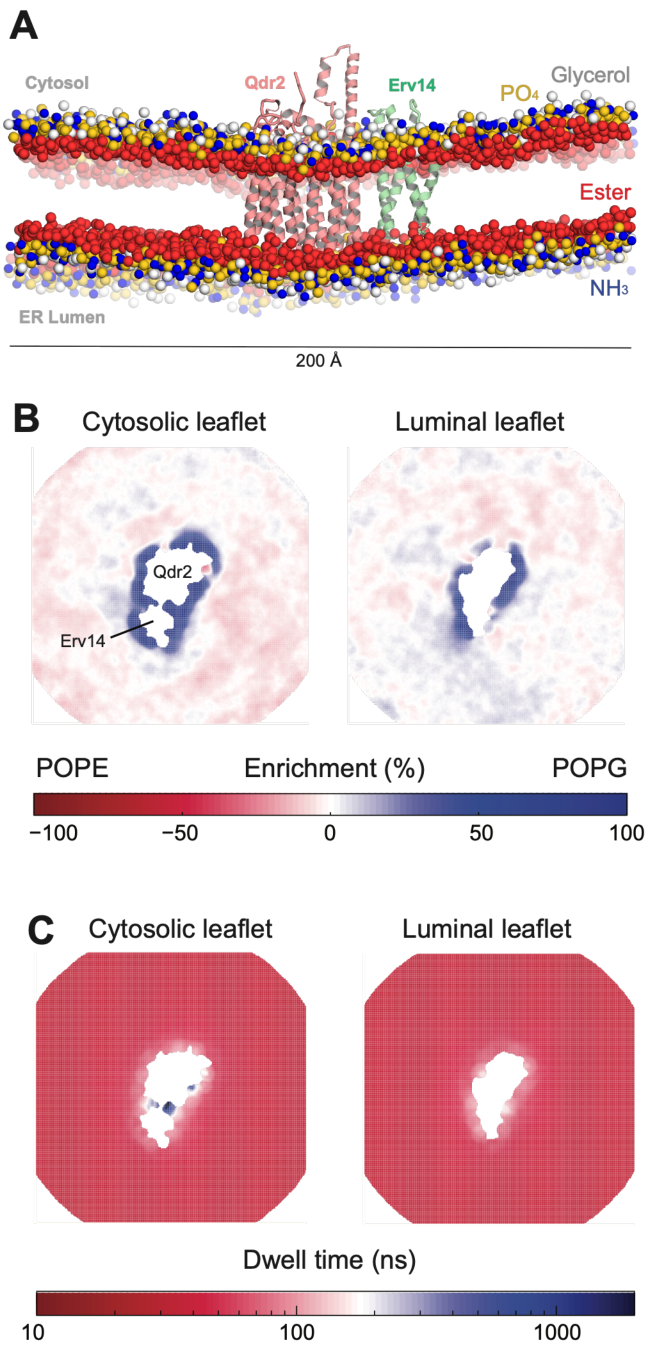


**Fig. S3. Molecular dynamics simulations of the Erv14-Qdr2 complex in a simplified membrane. (A)** Snapshot of the POPE/POPG bilayer containing the Erv14-Qdr2 complex after 20 μs of simulation, viewed in the membrane plane. Qdr2 (pink) and Erv14 (green) are shown as cartoons and were internally constrained during the simulation; lipids are rendered as spheres colored by chemical moiety: choline/glycerol (white), phosphate ($PO_4$, yellow), ester (red), and ammonium ($NH_3$, blue). The acyl tail, water and ion particles are hidden for clarity. The cytosol is at the top and the ER lumen at the bottom. Scale bar, 200 Å. **(B)** Time-averaged lipid enrichment maps for the cytosolic (left) and luminal (right) leaflets, projected onto the membrane plane. Color indicates local enrichment (%) of POPG (blue) versus POPE (red) relative to the bulk composition, according to the scale below. The protein-occupied region is masked in white; the positions of Qdr2 and Erv14 are indicated. POPG is enriched in the annular shell surrounding the complex, most prominently in the cytosolic leaflet. **(C)** Corresponding maps of lipid dwell time (ns) for the cytosolic (left) and luminal (right) leaflets, colored on a logarithmic scale as indicated below. Long-lived lipid interaction sites (blue) coincide with regions of POPG enrichment adjacent to the protein.

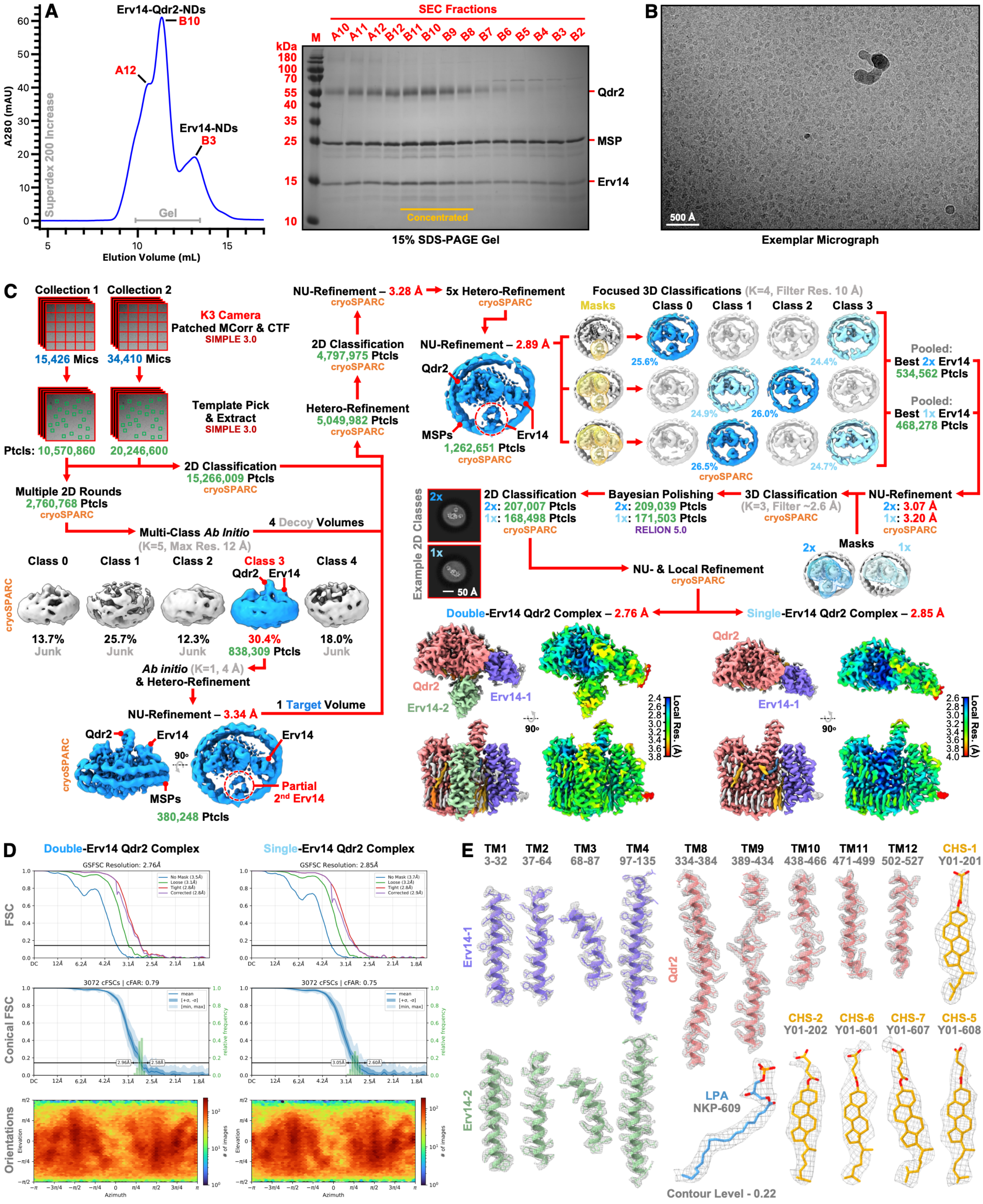


**Figure S4. Cryo-EM workflow, map resolution and model density fit for the Erv14-Qdr2 complexes in nanodiscs. (A)** SEC profile of the purified Erv14-Qdr2 complex reconstituted into nanodiscs with the peak corresponding to the Erv14-only nanodisc subpopulation labelled. The grey bar indicates the fractions shown in the Coomassie-stained SDS-PAGE gel, and the orange bar those fractions pooled and concentrated for cryo-EM. **(B)** Representative micrograph. **(C)** Overview of data acquisition and processing workflow with the software packages employed at each stage indicated. Local resolution estimates for the sharpened maps (contour level 0.3-0.34) are also shown. **(D)** Gold-standard Fourier shell correlation (FSC) curves estimating global resolution (0.143 cut-off) alongside conical FSC and orientation plots for the final reconstructions. **(E)** Density fits of the double-Erv14 complex for all transmembrane helices of Erv14-1 and Erv14-2, as well as select interfacial lipids and notable Qdr2 helices.

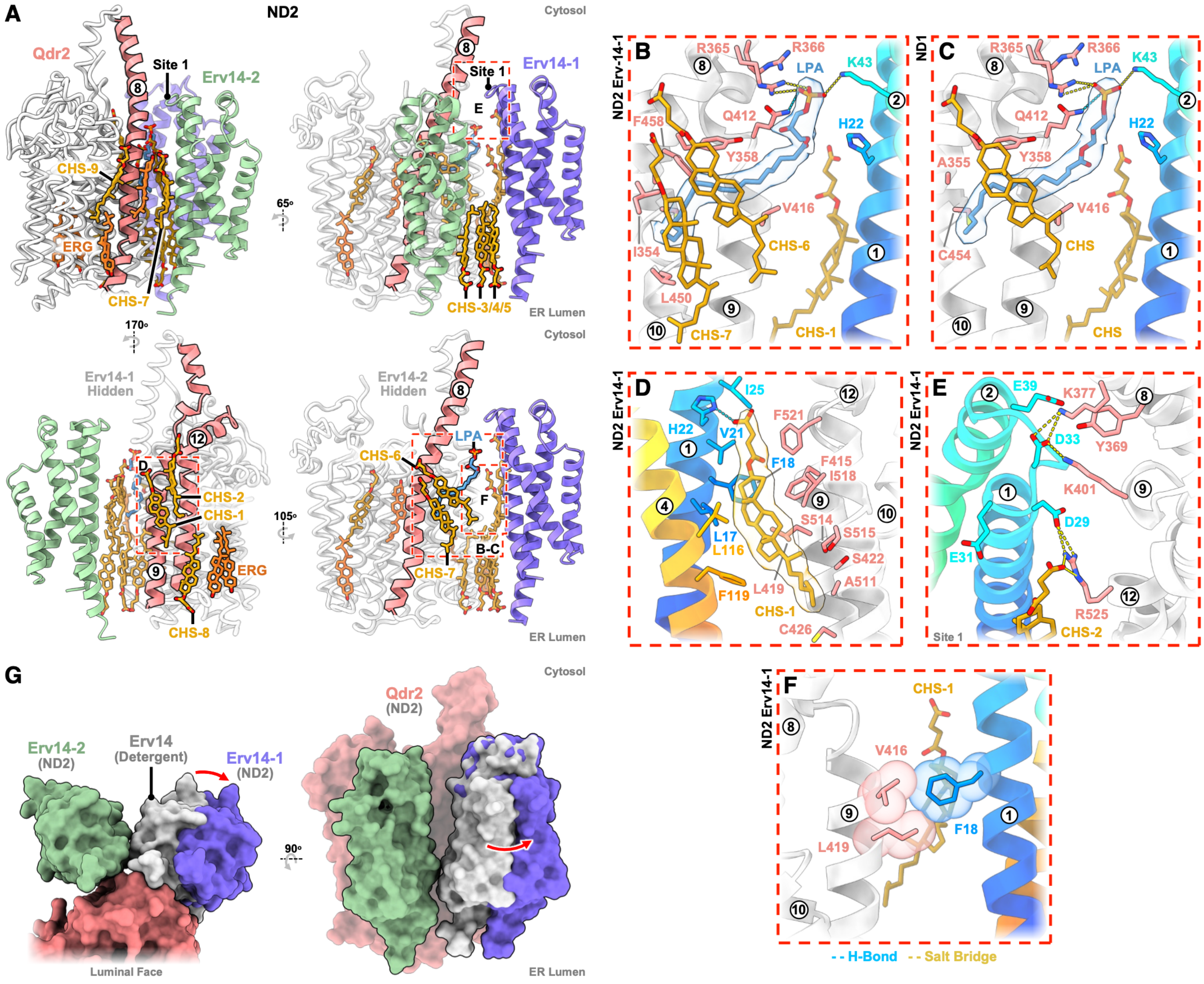

**Figure S5. Interactions between Erv14 and Qdr2 in nanodiscs. (A)** Views of the ND2 Erv14-Qdr2 complex highlighting all modelled CHS and ERG molecules as well as the resolved LPA. The two bottom views emphasise lipids populating the Erv14-1-Qdr2 or Erv14-2-Qdr2 interfaces that are otherwise obscured by the respective cargo receptor in the top two representations. Boxed regions correspond to the interaction close-ups shown in panels **(B-F)**, which are coloured as in the main text. Where shown, cryo-EM lipid density is contoured at 0.22 (sharpened map). Zoomed in views: The Erv14-1-Qdr2 LPA binding site in the **(B)** ND2 and **(C)** ND1 structures, **(D)** Interactions formed by CHS1 at the ND2 Erv14-1-Qdr2 interface, (**E)** Salt bridges between ND2 Erv14-1 Site 1 residues and Qdr2 and **(F)** The "knobs-in-holes" arrangement of ND2 Erv14-1 F18 and hydrophobic residues on Qdr2 TM9. **(G)** Surface representations of the Erv14-Qdr2 complex viewed from the membrane plane and the ER lumen, illustrating the relative rearrangement of Erv14-1 following lipid remodelling in nanodiscs.

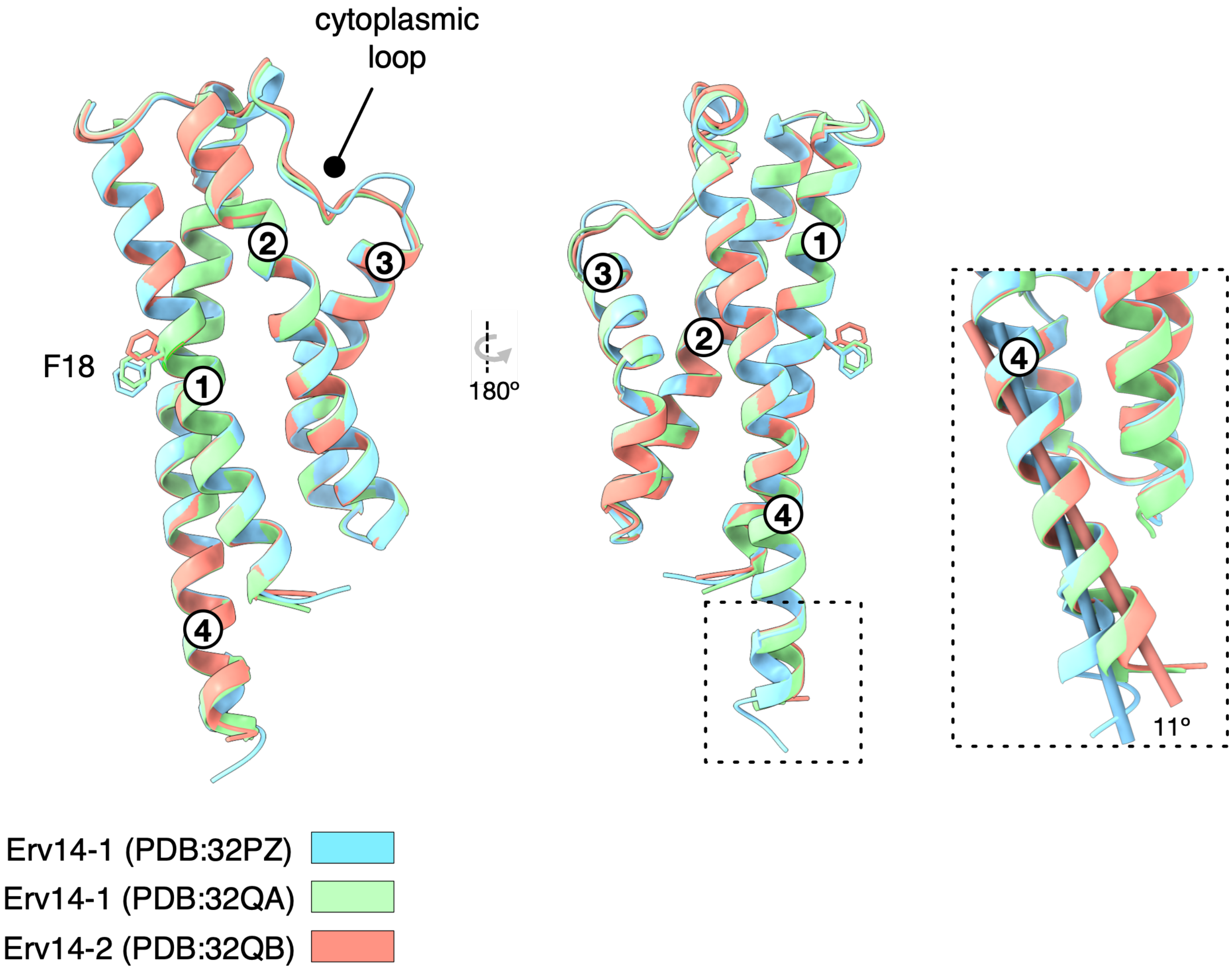


**Figure S6. Structural analysis of Erv14.** Overlay of the Erv14 structures determined both in detergent (PDB:32PZ) and nanodisc structures Erv14-1 (PDB:32QA) and Erv14-2 (PDB:32QB). There is a slight movement of the extreme C-terminus between the detergent and nanodisc structures of Erv14-1.

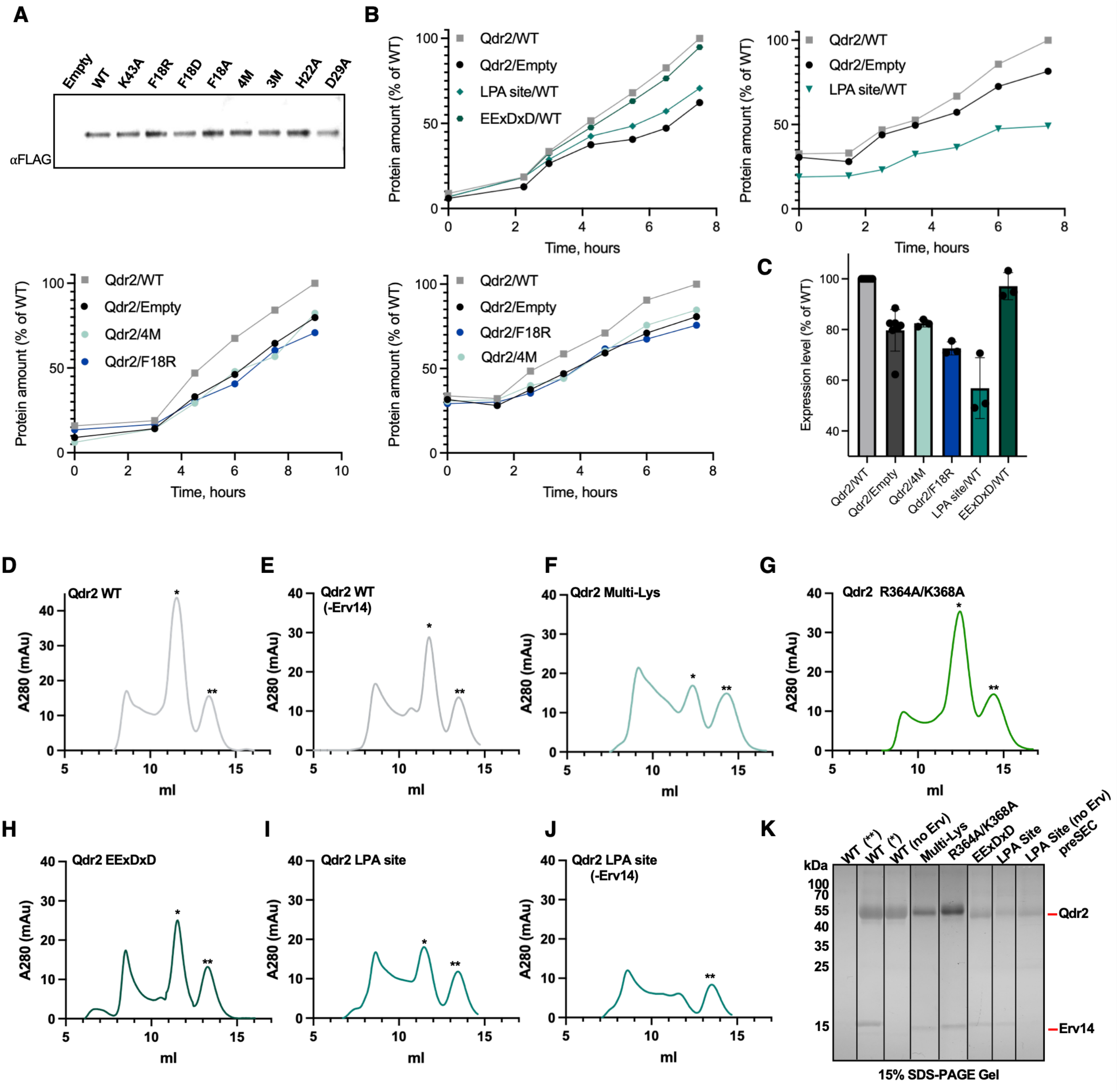


**Figure S7. Impact of Erv14 expression and engagement on Qdr2 biogenesis and stability. (A)** Western blot analysis of the Erv14-FLAG mutants used in the trafficking assay (Figure 4) indicates that the introduced mutations do not alter protein abundance in the cell. **(B)** Higher levels of Qdr2-GFP protein are produced in the presence of WT Erv14 (Qdr2/WT) compared with the absence of Erv14 (Qdr2/Empty – empty refers to the empty plasmid). Erv14 mutants that fail to interact with Qdr2, 4M (alanine mutations of H22, D29, E31 and D33) and F18R, do not increase the level of Qdr2. The lysolipid site (LPA site) mutations in Qdr2 result in reduced protein levels (and slower expression) due to protein instability and/or misfolding of these variants, as shown in panel D. The $E^{383}$ExDxD mutant on Qdr2 is synthesised at a similar level to WT. Representative data are shown for each mutant. **(C)** Quantification of Qdr2-GFP biogenesis data (shown in panel B). n=3, except for Qdr2/WT and Qdr2/Empty, where n=7, as these were repeated with every experiment. **(D-J)** Size-exclusion chromatography traces of purified Qdr2 protein, both WT **(D-E)** and mutants, purified in the presence of overexpressed Erv14 unless stated otherwise (endogenous Erv14 levels were present throughout). The Qdr2 Multi-Lys mutant **(F)** showed decreased stability, as indicated by protein extending from the void peak, suggesting increased aggregated protein. In addition, this mutation solubilised less well than the WT protein in LMNG: CHS. For WT, solubilisation efficiency was typically ~46-53%, and for this charged-site mutant (Multi-Lys) it was ~31-35%. **(G)** Mutation of the Erv14-2 interaction site on Qdr2 (R364A, K368A) results in stable, monodisperse protein. A similar result was observed for the EExDxD Sec24 binding mutant **(H)**. **(I)** The LPA site mutant reduced Qdr2 levels and stability (and demonstrated reduced solubility levels at ~28%). However, the stability of the LPA site mutant, as measured by its ability to be purified, is enhanced by Erv14 overexpression (**I** compared to **J**). **(K)** Coomassie-stained SDS-PAGE gel showing peak fractions for the Erv14:Qdr2 complex (*). The second peak observed in all chromatograms (**) contained no protein and is most likely excess detergent or lipid. For the LPA site mutation with no Erv14, the pre-SEC sample was loaded onto the gel because the Qdr2 peak was so low in this mutant. The Qdr2 Multi-Lys mutant, while showing some instability, still binds Erv14. A mutation at the Erv14-2 interaction site retains the ability to bind Erv14 when purified in LMNG: CHS. Given that this mutant is unable to traffic to the plasma membrane, this result suggests that the interaction with Erv14-1 is retained, as this is the only interaction we observe in this detergent in our structure. The $E^{383}$ExDxD site mutant also still interacts with Erv14, indicating its ability to interact with Erv14-1. Given the structural position of this mutation remote from the membrane, it is likely that it still interacts with Erv14-2.

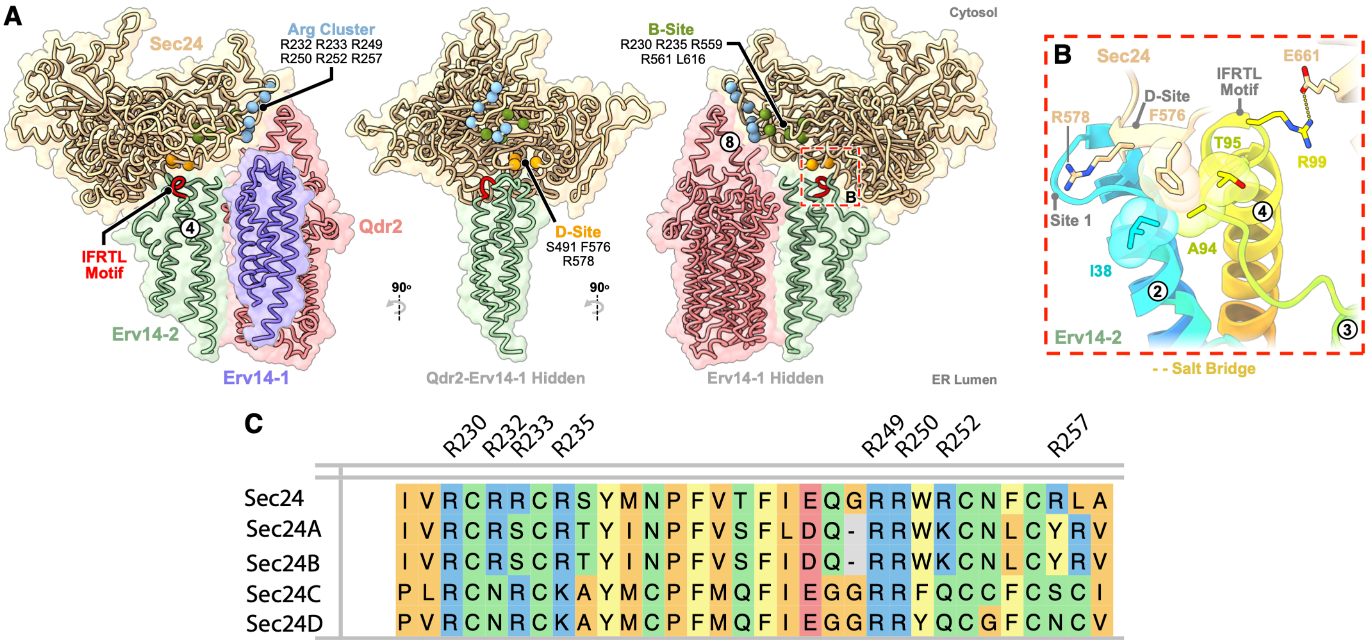


**Figure S8. Structural model for Sec24 recruitment during COPII coat formation. (A)** Views of the AlphaFold-generated model of the Erv14-Sec24 complex (shown in Figure 4G), highlighting the interaction site between Erv14-2 (near the IFRTL motif) and Site-D on Sec24 and the conserved arginine cluster on Sec24 with the extended TM8 of Qdr2. **(B)** Close-up of the interface between the region adjacent to the IFRTL motif on Erv14-2 with Site-D on Sec24 highlighting a nearby salt-bridge between R99 and E661. **(C)** Sequence alignment of the arginine cluster of Sec24 predicted to interact with Qdr2. Sequences shown are from *S. cerevisiae* (Sec24) and the human Sec24 paralogues (Sec24A-D).

**Table S1.** **Cryo-EM data collection, refinement and validation statistics.**

| | **Erv14-Qdr2 LMNG:CHS** (EMD-59064) (PDB 32PZ) | **Double-Erv14 Qdr2 NDs** (EMD-59065) (PDB 32QA) | **Single-Erv14 Qdr2 NDs** (EMD-59066) (PDB 32QB) |
|---|---|---|---|
| **Data Collection & Processing** | | | |
| Magnification | 105,000x | 105,000x | 105,000x |
| Voltage (kV) | 300 | 300 | 300 |
| Electron Exposure ($e^{-}$/Å$^2$) | 39.1 & 41.1 | 40 & 39.9 | 40 & 39.9 |
| Defocus Range ($\mu$m) | -2.2 to -1 | -2.2 to -1 | -2.2 to -1 |
| Pixel Size (Å) | 0.832 | 0.832 | 0.832 |
| Symmetry Imposed | C1 | C1 | C1 |
| Initial Particle Images (no.) | 21,293,856 | 30,817,460 | 30,817,460 |
| Final Particle Images (no.) | 373,696 | 207,007 | 168,498 |
| Map Resolution (Å) | 2.83 | 2.76 | 2.85 |
| FSC Threshold | 0.143 | 0.143 | 0.143 |
| Map Resolution Range (Å) | 2.50-14.54 | 2.40-32.01 | 2.60-38.23 |
| **Refinement** | | | |
| Initial Model Used (PDB code) | AlphaFold 3 | 32PZ | 32QA |
| Model Resolution (Å) | 3.0 | 2.9 | 3.1 |
| FSC Threshold | 0.5 | 0.5 | 0.5 |
| Map Sharpening $B$ Factor (Å$^2$) | -125.2 | -100.1 | -103.5 |
| Model Composition | | | |
| Non-hydrogen atoms | 5135 | 6505 | 5250 |
| Protein Residues | 627 | 764 | 628 |
| Ligands | 7 | 14 | 10 |
| $B$ Factors (Å$^2$) | | | |
| Protein | 67.35 | 68.85 | 75.11 |
| Ligand | 66.54 | 74.42 | 86.29 |
| R.M.S. Deviations | | | |
| Bond Lengths (Å) | 0.009 | 0.006 | 0.005 |
| Bond Angles (°) | 1.242 | 1.128 | 1.094 |
| Validation | | | |
| MolProbity Score | 0.98 | 0.88 | 0.95 |
| Clashscore | 2.09 | 1.42 | 1.85 |
| Poor Rotamers (%) | 0.57 | 0.31 | 0.38 |
| Ramachandran Plot | | | |
| Favored (%) | 98.56 | 98.68 | 98.88 |
| Allowed (%) | 1.44 | 1.32 | 1.12 |
| Disallowed (%) | 0 | 0 | 0 |

| Property | Erv14-Qdr2 (PDB: 32PZ) | Erv14-1-Qdr2 (PDB: 32QA) | Erv14-2-Qdr2 (PDB:32QA) | Interpretation |
|---|---|---|---|---|
| Buried interface area | 733 Å² | 401 Å² | 258 Å² | Progressive reduction in direct protein–protein packing |
| $\Delta G_{solv}$ P-value* | 0.132 / 0.077 | 0.954 / 0.843 | 0.678 / 0.675 | Transition from a predominantly hydrophobic interface to increasingly polar interfaces as lipids interdigitate between the receptor and the cargo |
| Hydrogen bonds | 3 | 5 | 2 | The ND2 Erv14 interface compensates for reduced surface area with an expanded hydrogen-bond network |
| Salt-bridge contacts | 3 | 6 | 2 | Electrostatic interactions become increasingly important in the ND structure |
| Conserved interaction | R525–D29 | R525–D29 retained and expanded | R364–D29 equivalent interaction | Conserved electrostatic anchor despite interface remodelling |
| LPA lipid site | Conserved | Conserved | Conserved | Lipid binding is maintained despite changes in protein packing |
| CHS sites | Two hydrophobic pockets | Multiple hydrophobic pockets | Multiple hydrophobic pockets | Lipid-binding environment is redistributed with the altered quaternary arrangement |

**Table S2. Manual annotation and PDBePISA analysis of the binding interfaces between Qdr2 and Erv14 using the LMNG: CHS and ND2 structures.** The LMNG: CHS complex (PDB:32PZ) is dominated by a single extensive hydrophobic interface. It buries 733 Å², has the lowest $\Delta G_{solv}$ P-values, and relies on only three hydrogen bonds and a single R364–D29 electrostatic anchor. The ND2 Erv14-1-Qdr2 packing represents a remodelled interface. Although the buried surface area is reduced by ~45%, it exhibits almost twice as many hydrogen bonds and a much larger electrostatic network. The accompanying increase in $\Delta G_{solv}$ P-values is entirely consistent with a shift from hydrophobic packing to a more polar interface. The ND1 Erv14-2-Qdr2 complex forms the smallest interface, but it is not simply a weak version of the Erv14-1 interface. Instead, it has a compact, well-defined interaction network centred on R364-D29, supplemented by a cation-pi interaction between Y34 and R364, and by a cation-pi interaction between K368 and R364 (Fig. 3B). * $\Delta G_{solv}$ P-value measures the probability that a randomly chosen surface patch of the exact same surface area on the protein would have a lower (more favourable) solvation free energy gain ($\Delta_i G$) than the observed interface (*19*).
